# Crossmodal representations in the auditory cortex of deaf and hearing individuals

**DOI:** 10.1101/2025.07.11.664439

**Authors:** Barbara Manini, Konstantin Grin, Martin Eimer, Samuel Evans, Luigi Tamè, Bencie Woll, Rita Bertani, Dilay Ercelik, Lucy Core, Yueming Gao, Matthew Longo, Valeria Vinogradova, Velia Cardin

## Abstract

The study of early deafness provides unique insights into how sensory experience shapes brain function and organisation. Here we investigated whether the auditory cortex of deaf individuals can implement cognitive functions when its sensory input is absent or significantly reduced. Crossmodal plasticity research has shown that auditory areas of the human brain are recruited for executive processing tasks in deaf individuals (Cardin et al., 2018; Ding et al., 2015; Manini et al., 2022; Zimmermann et al., 2021). What is the role of the auditory cortex during such executive processes in deaf individuals? One possibility is that it has a role in sensory processing, extracting information about relevant features. Alternatively, their function may go beyond feature processing, representing information such as task rules or attentional states. To distinguish between these hypotheses, we conducted an fMRI delay-to-match experiment in either the visual or somatosensory modality in deaf (N=13) and hearing (N=18) individuals. Representational Similarity Analysis (RSA) showed that the auditory cortex of deaf individuals contains information task and sensory modality. We also found significant representations of somatosensory frequency. Critically, task and modality representations were also found in the auditory cortex of hearing individuals. These findings suggest that crossmodal plasticity relies on the enhancement of representations that are present in hearing individuals, rather than through the implementation of novel ones. In conclusion, we show that sensory experience shapes cognitive processing and the function of sensory regions in the brain, and that the “functional destiny” of cortical regions can be shaped by early sensory experience.

## Introduction

Information from our senses is constantly used to learn and understand our environment. This sensory experience plays a crucial role in shaping the brain. Understanding this process is central in determining the extent to which the function of brain regions is predetermined from birth, and how much it is shaped by environmental experience. Here we ask whether brain regions typically considered ‘sensory’ can implement cognitive functions when their main input is absent or significantly reduced during development. We address this by studying the responses of the auditory cortex to visual and somatosensory cognitive tasks in congenitally and early deaf individuals, and evaluating the representation of information across different dimensions: stimuli features, task and sensory modality.

In the mammalian brain, sensory cortices have a prominent role in perceptual processing, receiving strong inputs from the sensory organs (Krubitzer, 1995). However, in individuals who are deaf or blind from birth, these inputs are absent or significantly reduced. In such cases, it has been found that these regions process other sensory information, a phenomenon known as crossmodal plasticity (Cecchetti et al., 2016; Heimler et al., 2015; Merabet & Pascual-Leone, 2010). Not only do these cortices respond to stimulation in different sensory modalities, but there is evidence suggesting a specialisation for cognitive processing in the sensory cortices of deaf and blind individuals (Bedny et al., 2015; Cardin et al., 2018; Ding et al., 2015; Kanjlia et al., 2021; Manini et al., 2022; Zimmermann et al., 2021). Specifically, research in deaf individuals has shown that regions of the superior temporal cortex, which are usually considered auditory-processing regions, are recruited for executive processing in the visual and somatosensory modalities (Cardin et al., 2018; Ding et al., 2015; Manini et al., 2022; Zimmermann et al., 2021). These are functions that are typically associated with fronto-parietal regions of the brain, and not sensory cortices.

What is the role of sensory regions during such executive processes in deaf individuals? Studies that suggest a specialisation for executive functions are usually fMRI studies where sensory properties are kept constant across conditions, but task demands are manipulated. In such cases, activations in auditory cortices have been observed during a visuo-spatial delayed recognition task (Ding et al., 2015), a visual two-back working memory task (Cardin et al., 2018), visual and somatosensory temporal sequence discrimination tasks (Zimmermann et al., 2021), and a switching task in the visual modality (Manini et al., 2022). However, not all executive function tasks activate the auditory cortex in deaf individuals. For example, in a study where we tested four executive function tasks including visuo-spatial working memory, switching, planning, and inhibition, we only found significant activations in the auditory cortex of deaf individuals during the switching task (Manini et al., 2022). The fact that not all executive function tasks, and not even all working memory tasks, activate auditory cortex in deaf individuals, has led to suggest that rather than a general role in cognitive control, auditory regions in deaf individuals have a role in updating task-sets or reallocation of attention (Manini et al., 2022).

Previous studies of crossmodal plasticity have not been able to elucidate the nature of the functional representation in auditory regions of deaf individuals — whether this cortex is implementing task-specific information, such as updating task-sets or attentional control, or enhancing the representation of sensory features due to task demands. This is because most fMRI studies showing specialisation for cognitive tasks in the auditory cortex of deaf individuals have used univariate analyses, where results show overall differences in activations between conditions and/or groups, but cannot provide information about the specific information represented in a cortical region. Thus, it is possible that increased activations during executive function tasks in deaf individuals reflect sensory processing functions, such as extracting information about relevant sensory features. Alternatively, the univariate changes may reflect processing of task-specific information, such as updating task-sets or attentional control. We have previously referred to these hypotheses as ‘functional-preservation’ and ‘functional-change’ (Cardin et al., 2020), and they have distinct implications for our understanding of the brain and the role of sensory experiences in shaping its function. In the first case, it would suggest that sensory regions would preserve a sensory processing role, independently of the sensory experience of the individual. The second case, instead, implies that as a result of different sensory experience, crossmodal plasticity changes the function of the cortical region.

To address the nature of the functional role of the auditory cortex of deaf individuals, we conducted an fMRI experiment in which participants had to implement different tasks in the visual and somatosensory modality, while the overall properties of the stimuli were held constant. The assumption is that for each task, participants will use a set of different cognitive functions that will allow them to implement the task rules and successfully achieve the goal (Sakai, 2008). We will refer to these as task effects. We used Representational Similarity Analysis (RSA) to determine what information is represented in the auditory cortex of deaf individuals. RSA quantifies the similarity between neural patterns in a given brain area for a set of stimuli varying in one or more dimensions (Kriegeskorte et al., 2008). Applying RSA, we can test different hypothetical models of *how* information is represented in the auditory cortex, disentangling the effect of stimuli features, task and sensory modality (Kriegeskorte et al., 2008), and determining whether information about one or more of these dimensions is represented in this area.

If the auditory cortex of deaf individuals has a role in extracting stimuli features, we should find information about the low-level properties of the stimuli, with the RSA analysis favouring models of sensory features. In addition, if the auditory cortex of deaf individuals has a role in implementing a set of cognitive functions that allow the successful completion of a task, we should find a representation of task information. Representation of information could also differ at a more abstract level, reflecting cognitive processes that are implemented across all tasks in a given sensory modality. In such a case, we should find information about the sensory modality of stimulation, even in cortices that do not receive direct inputs from the stimulated sensory organs.

In summary, our study identifies whether information about stimuli features, task, and sensory modality can be decoded from activity in the auditory cortex of deaf individuals. This will help us understand how sensory experience shapes cognitive processing and the function of sensory regions in the brain, and the extent to which the “functional destiny” of cortical regions can be changed by early sensory experience.

## Materials and Methods

### Participants

There were two groups of participants:

- 13 severely-to-profoundly deaf individuals who were deaf at birth, and are native signers of British Sign Language (BSL) [mean age=37.9, sem=4.2, range= 19 – 69; 9 female, 4 male] (Table 1).
- 18 hearing individuals, native or proficient speakers of English (including 4 of the authors; 6 participants knew a sign language to conversational or proficient level) [mean age=34.4, sem=3.2, range= 19-60; 9 female, 9 male].

**Table 1.**
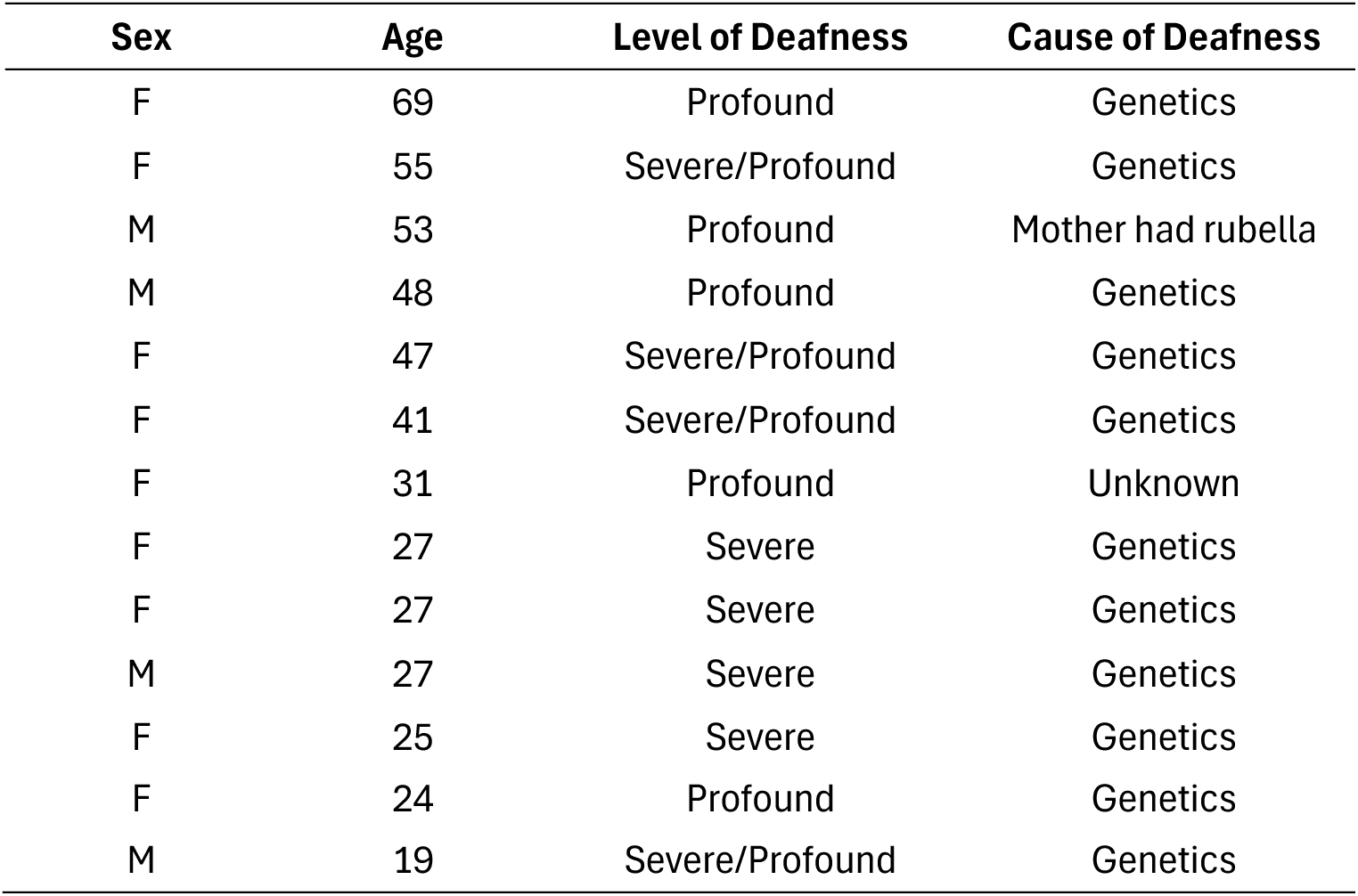
Demographics of deaf participants.

There was no significant difference in age between groups (t(29)=0.67, p=.51). All participants communicated with the researchers directly using BSL or English. All participants gave their written informed consent to take part in the study. All deaf participants were native BSL signers in order to avoid effects of potential delayed language acquisition on cognitive functions (Botting et al., 2017; Hall et al., 2017); they self-reported as severely or profoundly deaf.

All procedures followed the standards set by the Declaration of Helsinki and were approved by the UCL Research Ethics Committee. Participants travelled to University College London to take part in the study. They were recruited through public events, social media and the participant databases of the UCL Deafness, Cognition and Language Research Centre and UCL Psychology subjects pool (SONA). Participants were all right-handed (self-reported), had full or corrected vision, and no history of neurological conditions. All participants were compensated for their time, travel and accommodation expenses. One participant was scanned but excluded from the final sample due to incidental findings, two were excluded due to movement in the scanner and one was excluded due to an interruption of the scanning session.

### Scanning session

Participants were scanned at the Birkbeck-UCL Centre for Neuroimaging (BUCNI) in London using a 3.0 Tesla Siemens MAGNETOM Prisma scanner and a 32-channel head coil.

Functional data were obtained using a T2*-weighted multiband sequence (60 slices, TE=35.20ms, TR=1250 ms, FoV=212mm, MB acceleration factor = 4, Bandwidth = 2620 Hz/Px) with an in-plane resolution of 2×2mm. A structural T1-weighted scan with an in-plane resolution of 1×1mm was also acquired from each participant (MP-RAGE, 208 sagittal slices, FoV=256xmm, and slice thickness=1mm, TR 2300 ms, TE 2.98ms). Session-specific fieldmaps were also obtained to correct for magnetic field inhomogeneities (TE1=10ms, TE2=12.46ms, TR=1020 ms, FoV=192mm, slice thickness=2mm).

Participants took part in two or three scanning sessions, consisting of: 5-8 experimental runs, 1 anatomical scan, 1 fieldmap per session, 1 visual working memory localiser and 1 somatosensory working memory localiser. To improve task engagement and reduce experimental fatigue (Smith et al., 2018), 1 min videos of action films (‘Mission Impossible’ and ‘Spectre’) were shown after each experimental run.

### Main Experiment

Participants performed a delay-to-match working memory task in the visual and somatosensory modalities. Participants were asked to either remember the location of stimulation (spatial task) or remember the frequency of stimulation (temporal task). The stimuli always had both a spatial and a temporal component, but depending on the task, participants had to selectively attend and remember one of the stimuli features. See Figure 1 for a graphical description. Throughout the manuscript, we will refer to the following dimensions of the experiment:

**Figure 1.**
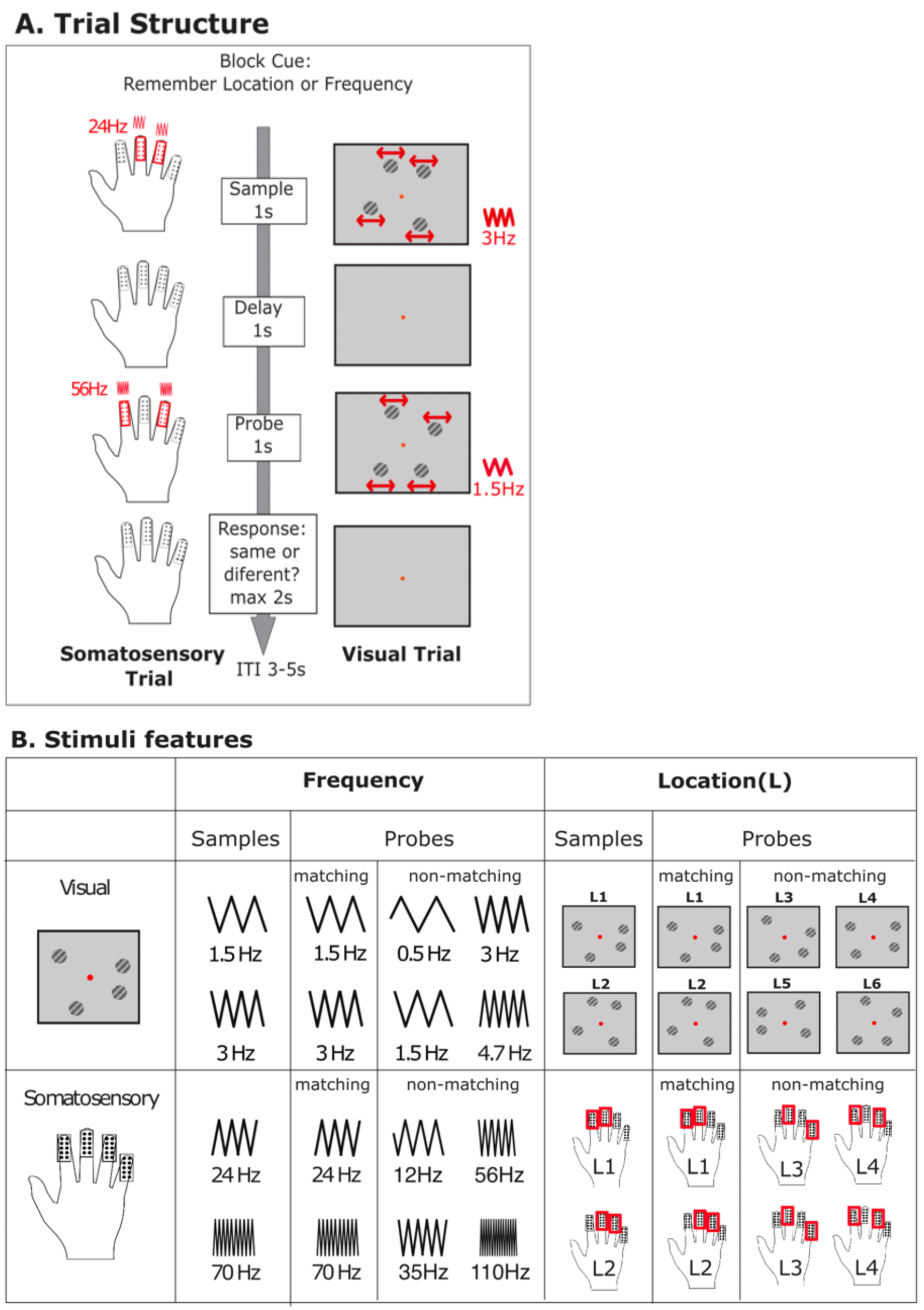
Experimental design and stimuli. Panel A: structure of an example trial in the visual and somatosensory modalities. Frequency of stimulation is shown in red. Red arrows in the visual trials indicate the lateral movement of the grating structure; these are for display purposes and were not present in the actual stimuli. Panel B: Stimuli features, showing the temporal frequency and spatial location of the visual and somatosensory stimuli. ITI: inter-trial interval. L: location. Red rectangles indicate the stimulated fingers.

- Stimuli features: Frequency and Location.
- Task: Spatial or Temporal
- Sensory Modality: Visual or Somatosensory

To optimise experimental design efficiency and statistical power, we prioritised increasing the number of trials over increasing the number of participants. The number of trials has a comparable effect on statistical efficiency to participant sample size (Chaumon et al., 2021; Chen et al., 2022; Nee, 2019). This approach is particularly advantageous for studies of special populations that are challenging to recruit, such as deaf native signers. Participants performed at least 5 and a maximum of 8 runs of the experiment across two or three scanning sessions. Each run lasted ∼12 min. Each run consisted of 2 blocks of each combination of sensory modality and task, for a total of eight blocks (2 visual temporal, 2 visual spatial, 2 somatosensory temporal and 2 somatosensory spatial). Before each block, participants saw a visual cue (2 sec) that indicated the task and sensory modality of the stimulation. Each block had 8 trials. During each trial, two stimuli were presented separated by a variable interstimulus interval (ISI). Throughout this paper, we will refer to the first stimulus of each trial as the ‘sample’ (presented for 1000ms); we refer to the 2^nd^ stimulus as the ‘probe’ (also presented for 1000ms). Sample and probe were separated by a variable ISI of 3000-5000ms (Fig 1). At the end of each trial, participants had 2 seconds to indicate whether the sample and probe matched in the cued feature. Participants responded with a right-handed button press, and the fixation dot turned from red to white to indicate that a response was recorded. After the participant’s response, there was a jittered intertrial interval (ITI) of 3000-5000ms. The main analyses was conducted on the sample stimuli (i.e. on the time window in which the sample was presented).

In each block (and across the experiment), 50% of the trials had matching samples and probes for the cued feature, and 50% had mismatching samples and probes. Half the probes were also a match for the non-cued feature. Values of the non-matching probes were chosen after several rounds of piloting to achieve performance of at least 75%. Stimuli were counterbalanced within and between runs to ensure equal temporal distance between unique samples across the experiment.

Before the scanning session, participants were familiarised with the tasks, and continued practising until they achieved at least 75% correct answers in all tasks.

### Visual stimuli

The visual stimuli consisted of four 3.3°- diameter circular gratings (eccentricity: 6.6°, grating-tilt: 30°) displayed on a grey background (Figure 1). The gratings were masked by a Gaussian mask. The texture of the gratings moved following a lateral uniform linear motion pattern, with the direction of motion changing at a frequency of 0.5-4.7 Hz. While the texture moved, the contour of the grating remained stationary, keeping the location of the whole grating constant. In the temporal task, participants had to remember the frequency at which the texture of the gratings changed its lateral direction of motion. Stimulus samples changed direction at either 1.5 or 3Hz. Probes changed direction at either 0.5, 1.5, 3 or 4.7Hz (Fig 1B). In the spatial task, participants had to remember the location of the stimuli. Visual location was defined as the spatial arrangement (polar angle) of the gratings on the screen. The samples had two patterns of locations, where gratings were positioned at the following polar angles: L1 — [155°, 19°, -24°, -117°] and L2 — [115°, 59°, -64°, -157°]. In the non-matching probes, two of the gratings were moved by ±20° in polar angle. This resulted in the following non-matching probes for L1: L3 — [135°, 19°, -44°, -117°], and L4 — [155°, 39°, -24°, -137°]. For L2 the nonmatching probes were: L5 — [135°, 59°, -44°, -157°] and L6 — [115°, 39°, -64°, - 137°] (Figure 1).

### Somatosensory stimuli

Somatosensory stimulation was delivered by a four-channel piezo-electric MR compatible stimulator (Quaerosys, Schotten, Germany, www.quaerosys.de). The stimuli consisted of tactile vibrations of skin indentation applied to the fingertips of the left hand. The stimulators were applied to the distal phalanges of the index, middle, ring, and little finger using Velcro tape, to ensure constant contact between the fingers and the stimulators throughout the experiment. The four stimulators in contact with the participant’s finger consisted of a matrix of 2 x 5 rods (1 mm in diameter), poking from a flat surface of 438 mm^2^. In the temporal task, participants had to remember the frequency of tactile stimulation. The frequency of stimulation of the sample stimuli was either 24Hz or 70Hz. For 24Hz samples, the stimulation frequency of the probes was either increased to 56 Hz or decreased to 12 Hz. For 70Hz samples, the stimulation frequency of the probes was either increased to 110Hz or decreased to 35Hz. In the spatial task, participants had to remember the location of stimulation. The location of the stimuli was determined by the stimulated fingers. In each event, two different fingers were stimulated at the same time (see Figure 1). The sample stimuli consisted of two patterns of location (L): L1 — index and middle finger; L2 — middle and ring finger. In the non-matching probes, we changed the stimulation of one of the fingers, resulting in the following patterns of stimulation: L3 — middle and little finger, or L4 — index and ring finger (Figure 1).

### Localisers

Independent visual and somatosensory working memory scans were used for the definition of the functional ROIs (see below). The visual localiser had a similar structure to the main delay-to-match working memory experiment, but trials were shorter: the sample and probe stimuli were presented for 500ms, the ISI was 1000-2000ms and the ITI was 1000ms. Participants also performed an additional condition where they had to monitor whether the stimuli changed in colour.

The somatosensory localiser also had a similar structure to the main experiment, but with shorter ISIs (1000-2000ms) and ITIs (2000-3000ms). There was an additional condition where participants had to monitor and identify the presence of a short gap in the somatosensory stimulation.

Each localiser had 3 blocks of each task, for a total of 9 blocks. For both localisers, the contrast [working memory > baseline] was used for ROI definition.

### ROI definition

Heschl’s gyrus (HG) and the superior temporal gyrus/superior temporal sulcus (STG/S) regions of interest (ROIs) were defined anatomically using FreeSurfer software (Fischl, 2012) (https://surger.nmr.mgh.harvard.edu) (Supplementary Fig 2 and 3). Each participant’s bias-corrected anatomical scan was parcellated and segmented (Dale et al., 1999; Fischl et al., 2002). The labels used to identify HG were: ‘ctx_lh_G_temp_sup-G_T_transv’ ID:11133 and ‘ctx_rh_G_temp_sup-G_T_transv’ ID:12133. The labels used to identify STG/S were *‘ctx_lh_G_temp_sup-Lateral’ ID:11134* and *‘ctx_rh_G_temp_sup-Lateral’ ID:12134*. Separately for each ROI, voxels with these were extracted and normalised to standard MNI space using SPM software (https://www.fil.ion.ucl.ac.uk/spm/).

The primary visual cortex (V1) ROI was defined using (Wang et al., 2015)’s probabilistic map combined with thresholded contrast images of the independent visual working memory localiser.

The primary somatosensory cortex (S1) ROI was defined using a probabilistic map of the representation of the left-hand digits in S1 (i.e., right S1; O’Neill et al., 2020), combined with thresholded images of the independent somatosensory working memory localiser.

Pre-supplementary motor area (preSMA) was chosen as a fronto-parietal control ROI because it is a multiple-demand region that responds to tasks in different sensory modalities and it is not biased towards a specific sensory modality (Noyce et al., 2017). preSMA was defined functionally, separately for each participant, using thresholded voxels from the contrast [working memory > baseline] from a combined analysis of the visual and somatosensory localisers. For each participant, we identified the peak activation voxel within a radius of 10mm around the coordinate [-2, 10, 50]. This voxel is the peak coordinate in preSMA when using the term ‘working memory’ in https://neurosynth.org/. After identifying each participant’s peak voxel, a volume of interest with a 10mm radius was defined using SPM.

In the definitions of V1, S1, and preSMA ROIs, localiser contrasts were thresholded at p < .005 (uncorrected). If this threshold resulted in ROIs with fewer than 95 voxels, p values were increased until at least 95 voxels were included in the ROI. In V1 and S1, voxels that were part of the probabilistic map and survived the significance threshold were included in the subject-specific ROI. See Supplementary Materials 1.

### Neuroimaging Data preprocessing

fMRI preprocessing was conducted using SPM12 (Wellcome Trust Centre for Neuroimaging, https://www.fil.ion.ucl.ac.uk/spm-statistical-parametric-mapping) implemented in MATLAB (MathWorks). The anatomical scans were segmented, bias-corrected, skull-stripped and normalised to the MNI (Montreal Neurological Institute) standard space.

The first 10 volumes of the functional data were discarded to account for T1 equilibrium effects. The remaining images were realigned to the first image, unwarped, coregistered to the skull-stripped anatomical image and normalised.

Univariate Neuroimaging Analysis.

Un-smoothed normalised images were used for the subject-specific first-level analysis, where a general linear model was estimated for all regressors. All the events were modelled as a boxcar and convolved with SPM’s canonical haemodynamic response function. Regressors were entered into a multiple regression analysis to generate parameter estimates for each regressor at every voxel.

The main regressors of interest were the stimulus sample presentations for the 4 main combinations of modality and task: visual temporal, visual spatial, somatosensory temporal and somatosensory spatial. Probes, cues and button presses were modelled separately as regressors of no interest. The motion parameters derived from the realignment of the images were also included in the model as covariates of no interest.

Results of the contrasts between the regressor of interest and the implicit baseline were extracted from each ROI and each participant using MarsBaR v.0.4440 (http://marsbar.sourceforge.net). The ROI analysis focused on the stimulus sample for consistency with the RSA analysis described below. The data were analysed using JASP (https://jasp-stats.org) and entered into separate mixed repeated measures ANOVAs. Factors for each ANOVA are reported in the relevant section of the Results.

### Representational Similarity Analysis (RSA)

RSA allows us to distinguish which features of a stimulus or task are represented in an observed neural response (Kriegeskorte et al., 2008). RSA calculates a Representational Dissimilarity Matrix (RDM), which quantifies the similarity (or dissimilarity by convention) between neural patterns in a given brain area for a set of stimuli varying in one or more dimensions (Kriegeskorte et al., 2008).

Our RSA analysis focused on the neural responses elicited by the sample stimuli. This was to achieve a large and equal number of condition-specific events and to avoid confounding effects from motor planning and execution. T-contrasts for each sample type and each run were estimated from unsmoothed subject-specific data. All other events (probes, cues, button presses) were modelled as regressors of no interest. For each participant and each run, we obtained 16 different T-contrasts (2 frequencies x 2 locations x 2 modalities x 2 tasks) representing all unique sample types (Fig 1B). This allowed us to test different models of representations of stimuli features, task and sensory modality (Figure 2). CoSMoMVPA (Oosterhof et al., 2016) was used to extract values from each subject specific T-contrast and ROI, removing non-finite and constant features and demeaning. Dissimilarity matrices for each ROI and each participant were obtained using RSA toolbox (Nili et al., 2014), by estimating the cross-validated Mahalanobis (crossnobis) distances using univariate noise normalisation (Walther et al., 2016). RDMs were exported to Matlab, and for each individual, the fit of different theoretical models (Fig 2) was calculated using CosmoMVPA by estimating the Pearson correlation coefficients between the theoretical model and each RDM, independently for each participant. Coefficients were then converting them into Fisher-transformed z-scores in Matlab.

**Figure 2.**
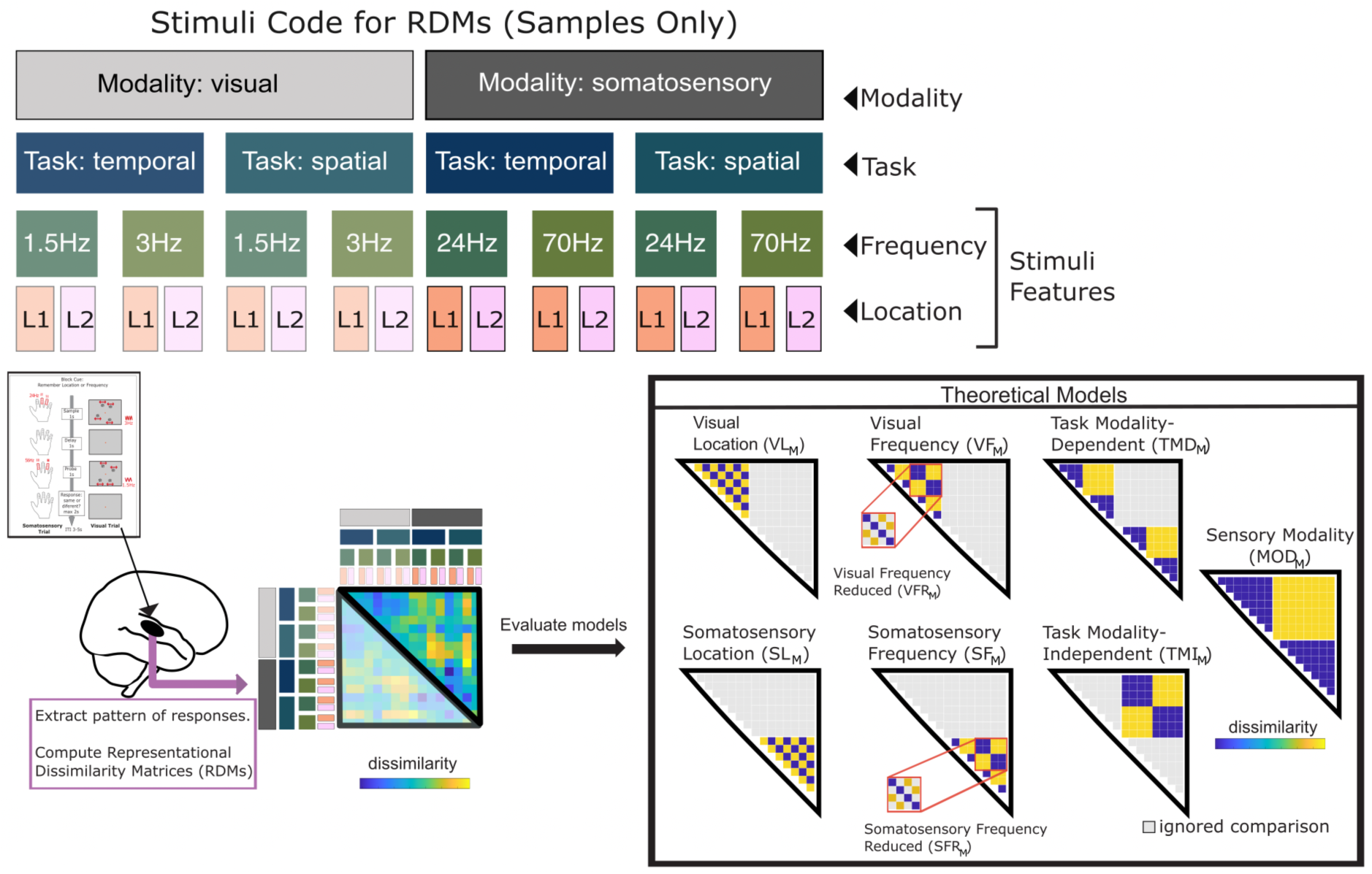
RSA analysis. The figure shows the structure of the Representational Dissimilarity Matrix (RDM) and a graphical representation of the theoretical models tested. There were three dimensions of interest: 1) Sensory Modality (Visual, Somatosensory); 2) Task (Temporal, Spatial) and 3) Stimuli features (Frequency, Location). L: location. See also Supplementary Figure 1.

The theoretical models represented different dimensions of the experiment. We tested the following stimuli-features models:

- Visual Location Model (VL_M_): codes the similarity in the spatial location of the visual stimuli.
- Visual Frequency Model (VF_M_): codes the similarity in the frequency of stimulation of the visual stimuli.
- Visual Frequency Reduced Model (VFR_M_): This model codes the similarity in the frequency of stimulation of the visual stimuli only for those samples with the same spatial location. In other words, it tests whether there is information about the frequency of stimulation when retinotopic stimulation is constant. Given the retinotopic organisation of the visual cortex (Engel et al., 1997), we considered it important to have a frequency model where the spatial location was consistent, in case retinotopy was preserved in the reorganised auditory cortex.
- Somatosensory Location Model (SL_M_): codes the similarity in the spatial location of the somatosensory stimuli.
- Somatosensory Frequency Model (SF_M_): codes the similarity in the frequency of stimulation of the somatosensory stimuli.
- Somatosensory Frequency Reduced Model (SFR_M_): This model codes the similarity in the frequency of stimulation of the somatosensory stimuli only for those samples with the same spatial location. Similarly to the VFR_M_ model for the visual modality, this model takes into account the somatotopic organisation of somatosensory cortices (Nelson & Chen, 2008; Penfield and Rassmussen, 1950), and considers that this might also be an organisational principle in reorganised auditory areas.

The following models tested task-based information:

- Task Modality-Dependent Model (TMD_M_): codes the similarity of the task within trials of the same sensory modality.
- Task Modality-Independent Model (TMI_M_): codes the similarity of the task across different modalities.

Finally, the Sensory Modality Model (MOD_M_) coded the sensory modality of stimulation across tasks.

When models tested the same section of the Representational Dissimilarity Matrix, we calculated the partial-correlation coefficient, to partial-out the contribution of the other models that varied in that section in space. This was the case for the VL_M_, VF_M_ and TMD_M_, and for the SL_M_, SF_M_ and TMD_M_ models.

To test for statistical significance, in control ROIs, we conducted a one-tailed t-test for each model to evaluate whether the Fisher-transformed correlation coefficients were significantly higher than zero. This results in three t-tests per model (one per control ROI). To correct for multiple comparisons, we applied a Bonferroni correction and set the significance threshold at p=.05/3=.017. If a model was a significant fit for a control ROI, we further conducted an independent-sample t-test to check for significant differences between groups, also using also a corrected p value of p=.05/3=.017.

In auditory ROIs we conducted repeated measures ANOVAs with the Fisher-transformed z-scores as the dependent variable, and the following factors: ROI (HG,STG/S), Hemisphere (right, left), and Group (deaf, hearing). Estimated marginal means obtained from JASP are reported to highlight those Fisher-transformed correlation coefficients that are significantly different from zero. To estimate the marginal means, JASP implements the R-package emmeans (https://cran.r-project.org/web/packages/emmeans/index.html), which estimates what the means would be if the design was balanced. When the assumption of sphericity was violated in the ANOVAs, a Greenhouse-Geisser correction was applied.

## RESULTS

### Behavioural Results

Figure 3 shows the behavioural results from the experiment. We conducted a 2 x 2 x 2 repeated measures ANOVA to evaluate whether there were significant differences in performance between sensory modalities, tasks and groups. Factors in the ANOVA included Modality (visual, somatosensory), Task (spatial, temporal), and between-subjects factor Group (deaf, hearing). There was a significant main effect of Task (F(1,29)=65.03, p<.001, ηp^2^=.69), driven by higher scores in the spatial tasks than in the temporal tasks ([spatial – temporal] = 0.071, SE=0.009, 95% CI=[0.053, 0.089]). There were no significant differences in performance between groups, with no main effect of Group and no interactions with Group. There was no main effect of Modality, and no significant interactions (but notice that the Modality * Task interaction values approach significance: F(1,29)=4.1, p=.052, ηp^2^=.12) (see full [Behavioural ANOVA here](https://osf.io/zqwt2)).

**Figure 3.**
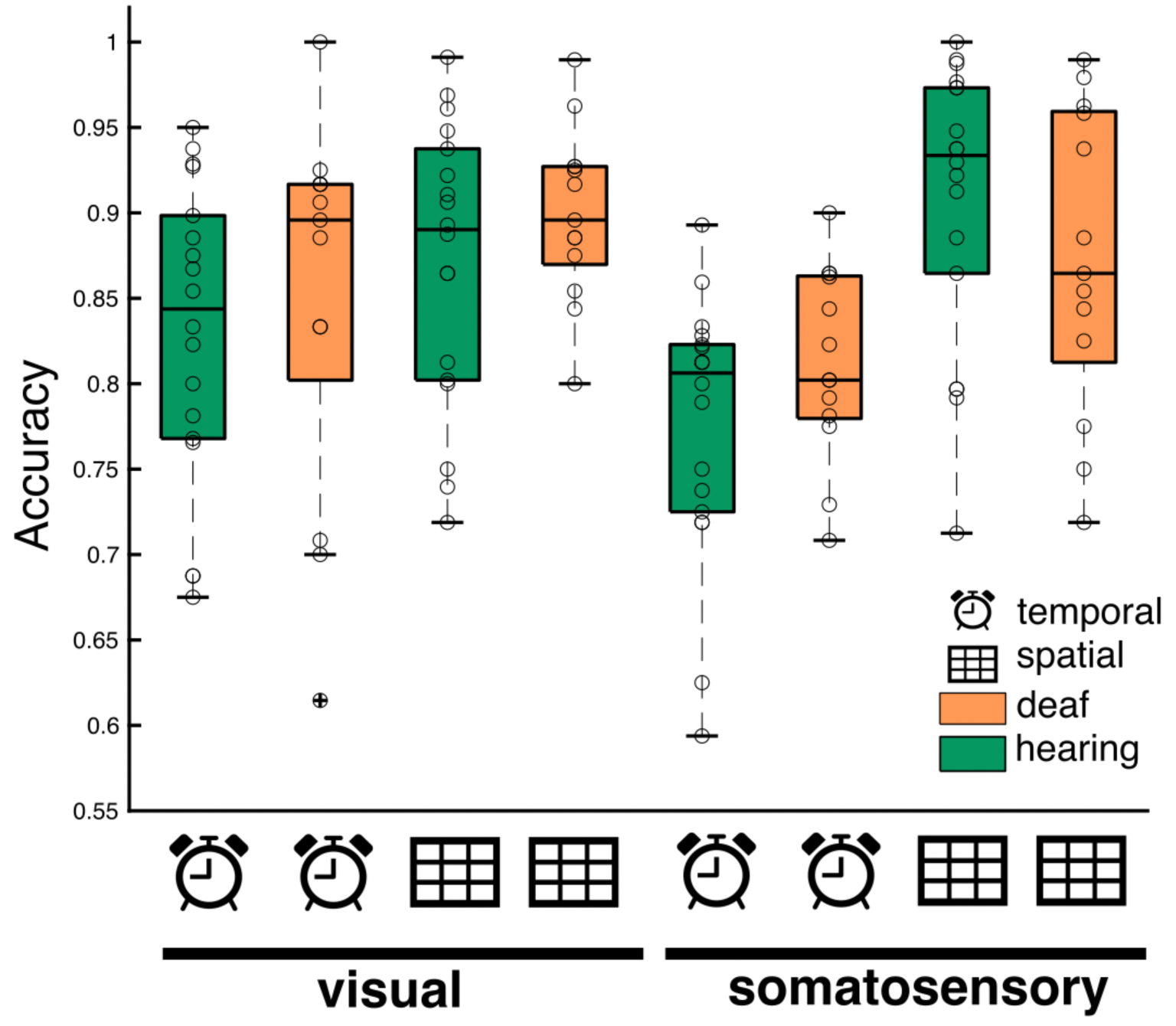
Behavioural results. Participants’ accuracy in the delay-to-match experiment averaged separately across groups, modalities and tasks. On each box, the edges indicate the 25^th^ and 75^th^ percentiles, and the black horizontal line inside the box indicates the median. See the [Behavioural ANOVA here](https://osf.io/zqwt2).

### Neuroimaging Results from Control ROIs

#### Univariate Analysis

We analysed three control ROIs: V1, where we expected responses to visual conditions; S1, where we expected responses to somatosensory conditions; and preSMA, where we expected significant responses to all conditions.

To evaluate this, we conducted repeated measures ANOVAs for each ROI separately, with contrast values from the univariate analysis as the dependent variable and the following factors: Modality (visual, somatosensory), Task (spatial, temporal), and Group (deaf, hearing).

Results from these ANOVAs showed the expected pattern of results (Figure 4; see the full [ANOVAs here](https://osf.io/9j5da)):

1. Significant activations for the visual conditions in V1, but not for the somatosensory conditions (Main effect of Modality: F(1,29)=68.8, p<.001, ηp^2^=.70; no significant interactions, see [V1 ANOVA](https://osf.io/9zpdt)).
2. Opposite pattern in S1, with significantly higher activations for the somatosensory conditions (Main effect of Modality: F(1,29)=137.06, p<.001, ηp^2^=.83; no significant interactions, see [S1 ANOVA](https://osf.io/zkdux)).
3. In preSMA, all visual and somatosensory conditions significantly activated this ROI (all marginal means significantly different from zero, p <.001; see [preSMA ANOVA](https://osf.io/35vuj)). There was a stronger response for temporal than spatial tasks, particularly in the somatosensory modality (Main effect of Task: F(1,29)=16.3, p<.001, ηp^2^=.36; Modality * Task interaction: F(1,29)=9.5, p=.004, ηp^2^=.25).
4. No significant main effects or significant interactions with Group in any of the control ROIs.

**Figure 4.**
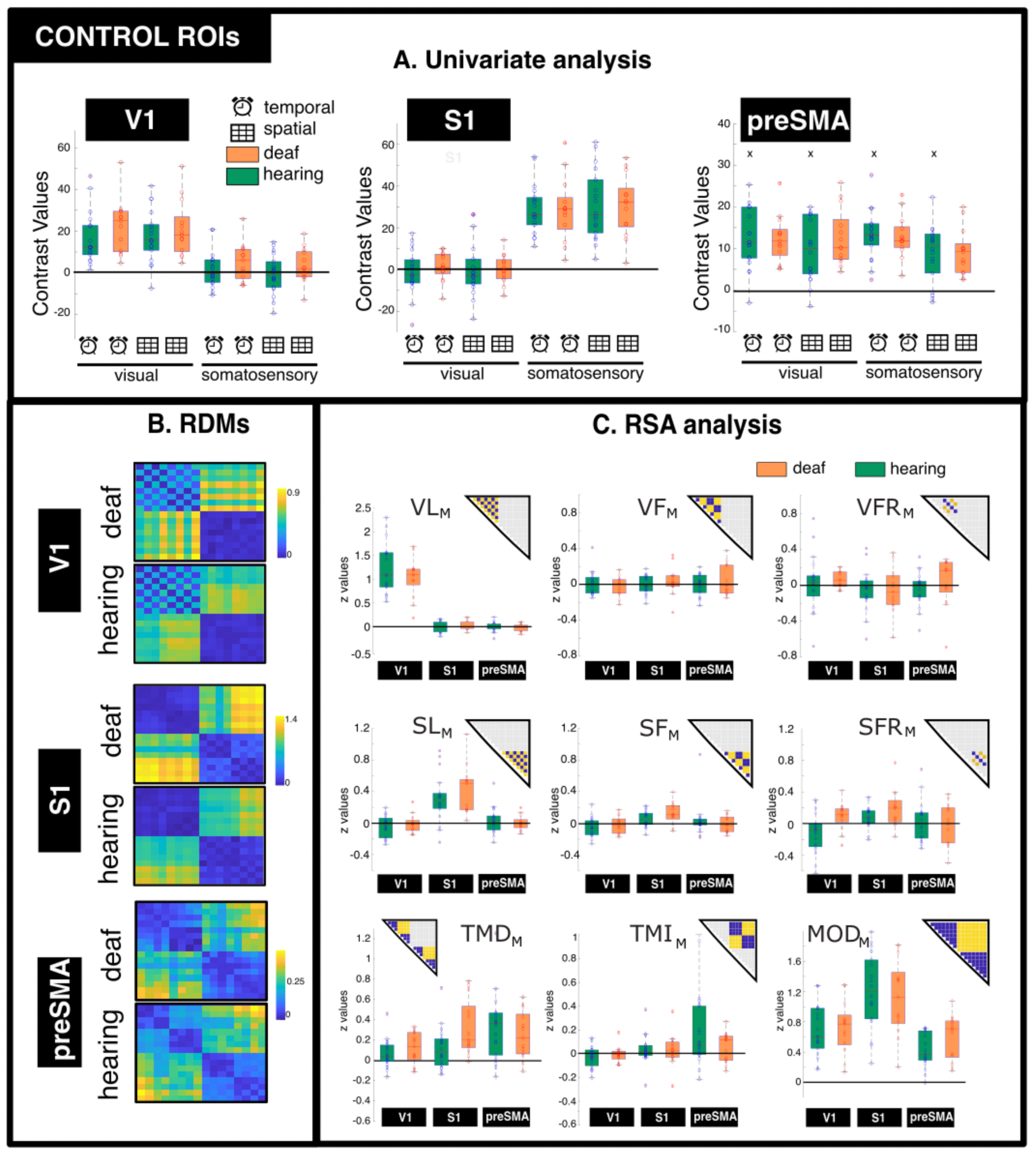
Results from Control ROIs. A: Univariate analysis results. Each data point represents the contrast values for a participant for a given condition vs the implicit baseline. x=outlier not shown in the graph. preSMA: pre-supplementary motor area. **B: Representational Dissimilarity Matrices (RDMs) for each group and each ROI.** The colour bar represents the value of the group-average cross-validated Mahalanobis (crossnobis) distance. Notice that the colour scale was set to the maximum and minimum value of each ROI to highlight within ROI differences across conditions, rather than highlighting between ROI differences which are not informative in our analysis. **C: Fisher-transformed z values for the Pearson correlation coefficients of relevant theoretical models.** The triangles above the the bar plots show the regions of the dissimilarity matrix that were evaluated in each model. VL**_M_**: Visual Location Model. VF**_M_**: Visual Frequency Model. VFR**_M_**: Visual Frequency Reduced Model. SL**_M_**: Somatosensory Location Model. SF**_M_**: Somatosensory Frequency Model. SFR**_M_**: Somatosensory Frequency Reduced Model. TMD**_M_**: Task Modality-Dependent Model. TMI**_M_**: Task Modality-Independent Model. MOD**_M_**: Sensory Modality Model. In the boxplots, the edges of each box indicate the 25^th^ and 75^th^ percentiles, and the black horizontal line inside the box indicates the median.

These results demonstrate the expected modality-specific responses in primary visual (V1) and somatosensory (S1) cortices, and common activations across modalities and tasks in the preSMA.

#### RSA analysis

In control ROIs, we expected the RSA analysis to reveal representations of stimuli features in sensory cortices (V1 and S1), and task information in preSMA. RDMs for the control ROIs are shown in Fig 4, and the statistical significance of the Fisher-transformed z-scores for each model can be found in Table 2. The RSA analysis confirmed our predictions (Fig 4).

**Table 2.** One Sample T-Tests for RSA Results in Control ROIs.

| ROIs | VL<sub>M</sub><br>T<sub>(30)</sub>, p | VF<sub>M</sub><br>T<sub>(30)</sub>, p | VFR<sub>M</sub><br>T<sub>(30)</sub>, p | SL<sub>M</sub><br>T<sub>(30)</sub>, p | SF<sub>M</sub><br>T<sub>(30)</sub>, p | SFR<sub>M</sub><br>T<sub>(30)</sub>, p | TMD<sub>M</sub><br>T<sub>(30)</sub>, p | TMI<sub>M</sub><br>T<sub>(30)</sub>, p | MOD<sub>M</sub><br>T<sub>(30)</sub>, p |
| --- | --- | --- | --- | --- | --- | --- | --- | --- | --- |
| V1 | **12.93, < .001** | -0.14, 0.550 | 0.96, 0.17 | -1.22, 0.88 | -1.633, 0.94 | -0.66, 0.743 | **3.38, .001** | -1.32, 0.902 | **11.5 < .001** |
| S1 | 0.19, 0.42 | 0.54, 0.30 | -1.3, 0.90 | **6.49, < .001** | **3.81, < .001** | **3.62, < .001** | **4.46, < .001** | 1.41, 0.084 | **13.1, < .001** |
| preSMA | -0.71, 0.76 | 0.51, 0.31 | -0.17, 0.56 | 0.89, 0.19 | 0.46, 0.33 | 0.32, 0.373 | **5.54, < .001** | **2.87, 0.004** | **10.2, < .001** |

The table shows results from one Sample T-Tests in Control ROIs for both groups combined. Significant $p$ values are highlighted in bold letters (Bonferroni corrected $p=.05/3=.017$ ). preSMA: pre-supplementary motor area. VL<sub>M</sub>: Visual Location Model. VF<sub>M</sub>: Visual Frequency Model. VFR<sub>M</sub>: Visual Frequency Reduced Model. SL<sub>M</sub>: Somatosensory Location Model. SF<sub>M</sub>: Somatosensory Frequency Model. SFR<sub>M</sub>: Somatosensory Frequency Reduced Model. TMD<sub>M</sub>: Task Modality-Dependent Model. TMI<sub>M</sub>: Task Modality-Independent Model. MOD<sub>M</sub>: Sensory Modality Model.

The z-scores for the Visual Location model (VL_M_) were significantly higher than zero in V1 (p <.001, Table 2). However, the Visual Frequency model (VF_M_) was not a good fit for this ROI or any of the control ROIs (p > .3). The z-scores for the Somatosensory Location model (SL_M_), Somatosensory Frequency model (SF_M_), and Somatosensory Frequency Reduced model (SFR_M_), were significantly higher than zero in S1 (p <.001, Table 2). In preSMA, the z-scores from task models were significantly higher than zero, both for the Task Modality-Dependent (TMD_M_, p <.001, Table 2; Fig 4) and the Task Modality-Independent (TMI_M_, p=.004, Table 2; Fig 4) models.

In addition, the z-scores for the Task Modality-Dependent model (TMD_M_) and the Sensory Modality (MOD_M_) model were significantly higher than zero in all control ROIs (Table 2; Fig 4). For all models and ROIs where the z-scores were significantly higher than zero, we conducted independent-sample t-tests to test if there were any significant differences between groups. None of the comparisons survived a Bonferroni-corrected p value of p<.017 (but notice an uncorrected significant difference in S1 for the task model; [Control ROIs Independent t-tests table](https://osf.io/et2un/files/bm89h)).

These results confirmed the expected pattern of results for control ROIs, with visual location represented in V1, and somatosensory location and frequency represented in S1. There was no significant representation of visual frequency in any of the control ROIs. In addition, task information was represented in a modality-dependent and independent way in preSMA. Task and sensory modality representations were also found in all control ROIs.

### Neuroimaging Results from Auditory ROIs

#### Univariate Analysis (Figure 5)

To identify crossmodal plasticity effects in deaf individuals, we analysed the univariate responses of four auditory ROIs: right Heschl’s Gyrus (RHG), left Heschl’s Gyrus (LHG), right superior temporal gyrus and sulcus (RSTG/S) and left superior temporal gyrus and sulcus (LSTG/S). To test for significant differences between groups we ran repeated measures ANOVAs for each ROI separately, with contrast values as the dependent variable and the following factors: Modality (visual, somatosensory), Task (spatial, temporal), and Group (deaf, hearing). Full results for the ANOVAs can be found here: [Auditory ROIs ANOVAs](https://osf.io/et2un/files/j863n).

**Figure 5.**
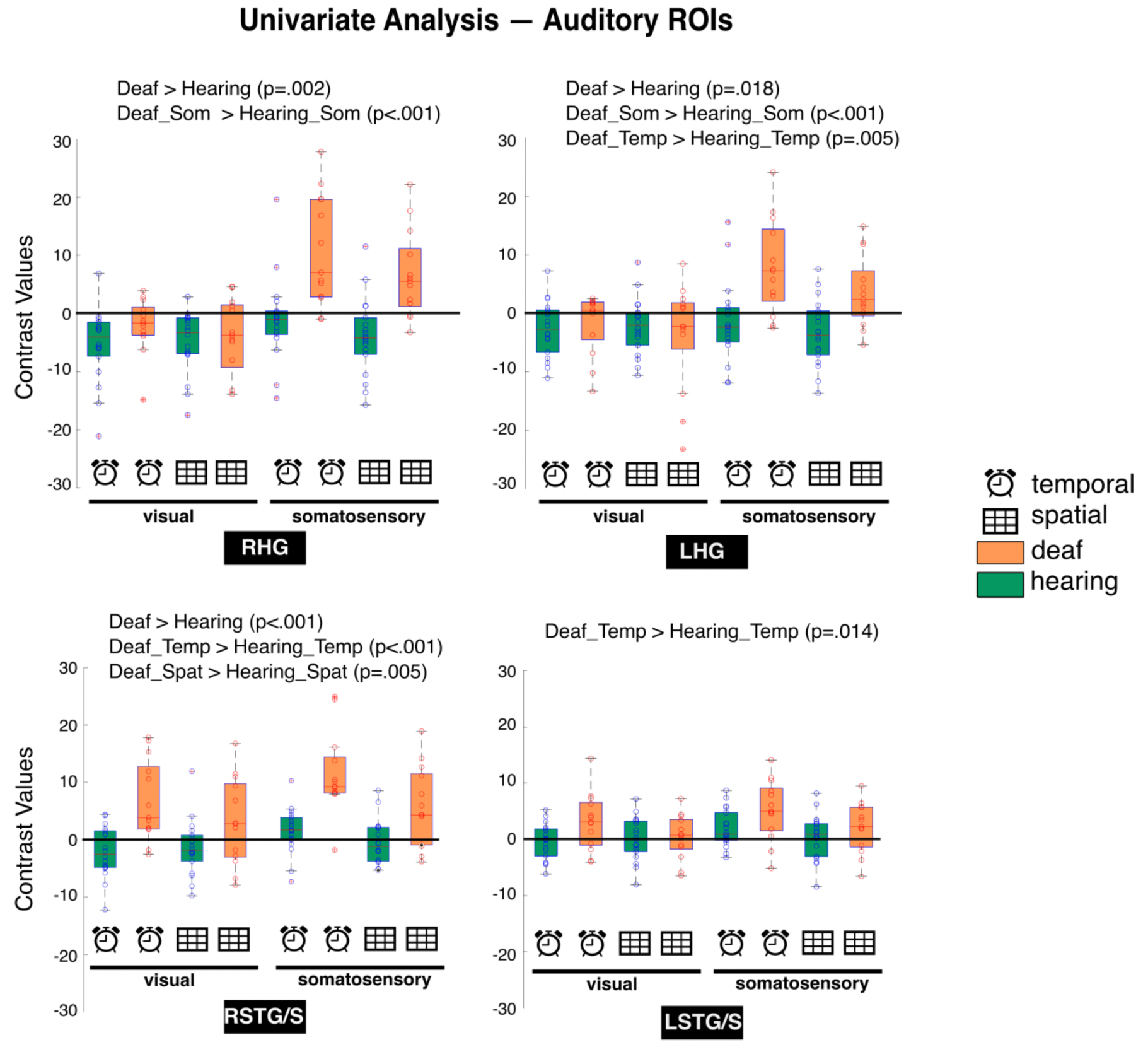
Univariate results for auditory Regions of Interest (ROIs). Each data point represents participant-specific contrast values for a given condition vs the implicit baseline. On each box, the edges indicate the 25^th^ and 75^th^ percentiles, and the black horizontal line inside the box indicates the median. Results for the comparison of interest are indicated above each panel (see [Supplementary Materials 2](https://osf.io/et2un/files/cjnke)). Full ANOVA results can be found here: [Auditory ROIs ANOVAs](https://osf.io/et2un/files/j863n). Temp: Temporal. Som: Somatosensory. Spat: Spatial. RHG: right Heschl’s gyrus; LHG: left Heschl’s gyrus; RSTG/S: right superior temporal gyrus and sulcus; LSTG/S: left superior temporal gyrus and sulcus.

Results from both RHG and LHG showed a significant main effect of Group (RHG: F(1,29)=11.94, p=.002, ηp^2^=.29; see [right HG ANOVA](https://osf.io/et2un/files/ugswv); LHG: F(1,29)=6.33, p=.018, ηp^2^=.18; see [left HG ANOVA](https://osf.io/et2un/files/eb7g9)). There was also a significant interaction Modality * Group in both right and left HG (RHG: F(1,29)=9.83, p=.004, ηp^2^=.25; LHG: F(1,29)=8.08, p=.008, ηp^2^=.22), reflecting increased responses in the somatosensory condition in the deaf group (Fig 5; see [Supp Mat 2 left HG](https://osf.io/et2un/files/768gz)). In left HG, there was also a significant Task * Group effect (F(1,29)=4.44, p=.044, ηp^2^=.13), reflecting significant differences between groups in the temporal task (Fig 5; [Supp Mat 2](https://osf.io/et2un/files/cjnke)).

In right STG/S, there was a significant main effect of Group (F(1,29)=19.1, p<.001, ηp^2^=.40), reflecting overall stronger responses in deaf individuals (Fig 5; see [right STG/S ANOVA](https://osf.io/et2un/files/t8k7e)). There was also a Task * Group interaction (F(1,29)=10.1, p=.003, ηp^2^=.26), with stronger responses in the temporal tasks in deaf individuals (Fig 5 and [Supp Mat 2 right STG/S](https://osf.io/et2un/files/qjnx3)).

In left STG/S, there was a significant Task * Group interaction (F(1,29)=5.7, p=.024, ηp^2^=0.16), reflecting an increased average response to the temporal tasks in the deaf group (Fig 5; see [left STG/S ANOVA](https://osf.io/et2un/files/msqa5) and [Supp Mat 2 left STG/S](https://osf.io/et2un/files/xrfas)). There was no significant main effect of Group (F(1,29)=4.0, p=.055, ηp^2^=0.12), and no significant Modality * Group interaction (F(1,29)=0.14, p=.71, ηp^2^=0.005).

In summary, results from the univariate analysis revealed crossmodal plasticity in the visual and somatosensory modality in auditory regions of deaf individuals. In HG, there was a clear stronger response to the somatosensory over the visual modality. In STG/S, there were significant responses in the visual and somatosensory modalities in deaf individuals, with stronger responses for the temporal over the spatial tasks.

#### Representational Similarity Analysis (RSA)

Figure 6 shows the RDMs for auditory ROIs. We conducted the RSA analysis to understand whether auditory regions in deaf individuals contain information about the different dimensions of our experiment (stimuli features, task and modality). To achieve this aim, we tested theoretical models that evaluated the different dimensions of the dataset (Fig 2). We were interested in understanding whether a given theoretical model was a significant fit for the data in a given ROI, and whether the representation of information was different between deaf and hearing individuals (rather than identifying which is the best theoretical model for a given ROI). This approach allows us to determine whether multiple representations of different dimensions coexist in the auditory cortex of deaf individuals. For that reason, for each model, we conducted an ANOVA with factors: ROI (HG, STG/S), Hemisphere (right, left), and Group (deaf, hearing). We report significant main effects and interactions, as well as significant marginal means, to highlight Fisher-transformed correlation coefficients that are significantly greater than zero. Full results for the ANOVAs can be found here: [Auditory ROIs RSA ANOVAs](https://osf.io/et2un/files/huvdq).

**Figure 6.**
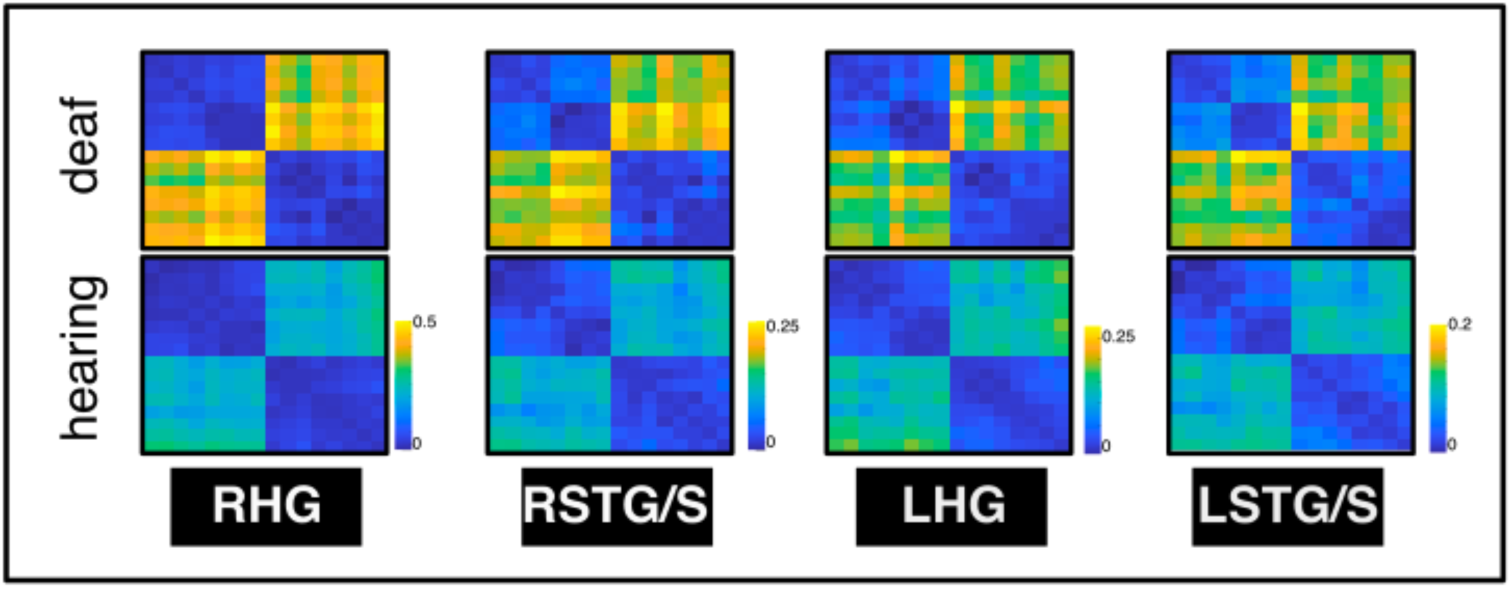
Representational Dissimilarity Matrices (RDMs) for each group and each auditory ROI. RHG: right Heschl’s gyrus; LHG: left Heschl’s gyrus; RSTG/S: right superior temporal gyrus and sulcus; LSTG/S: left superior temporal gyrus and sulcus. The colour bar represents the value of the group-average cross-validated Mahalanobis (crossnobis) distance. Notice that the colour scale was set to the maximum and minimum value of each ROI to highlight within ROI differences across conditions, rather than highlighting between ROI differences which are not informative in our analysis.

##### Representation of stimuli-features

Six of our theoretical models coded stimuli-features (Fig 2): Visual Location model (VL_M_), Visual Frequency model (VF_M_), Visual Frequency Reduced model (VFR_M_), Visual Frequency Reduced model (VFR_M_), Somatosensory Location model (SL_M_), Somatosensory Frequency model (SF_M_), and Somatosensory Frequency Reduced model (SFR_M_).

Of these models, the only one where we found significant differences between groups in auditory ROIs was the SFR_M_ [significant main effect of Group (F (1,29)=4.6, p=.04) (Fig 7; [SFR_M_ ANOVA](https://osf.io/et2un/files/pk85y))]. Marginal means for the deaf group were also significantly greater than zero (Deaf: mean=0.19, 95% CI=[0.07, 0.32], t(29)=3.2, p=.003; Hearing: mean=0.01, 95% CI=[-0.11, 0.13], t(29)=0.17, p=.86). There were no other significant main effects and interactions for this model.

**Figure 7.**
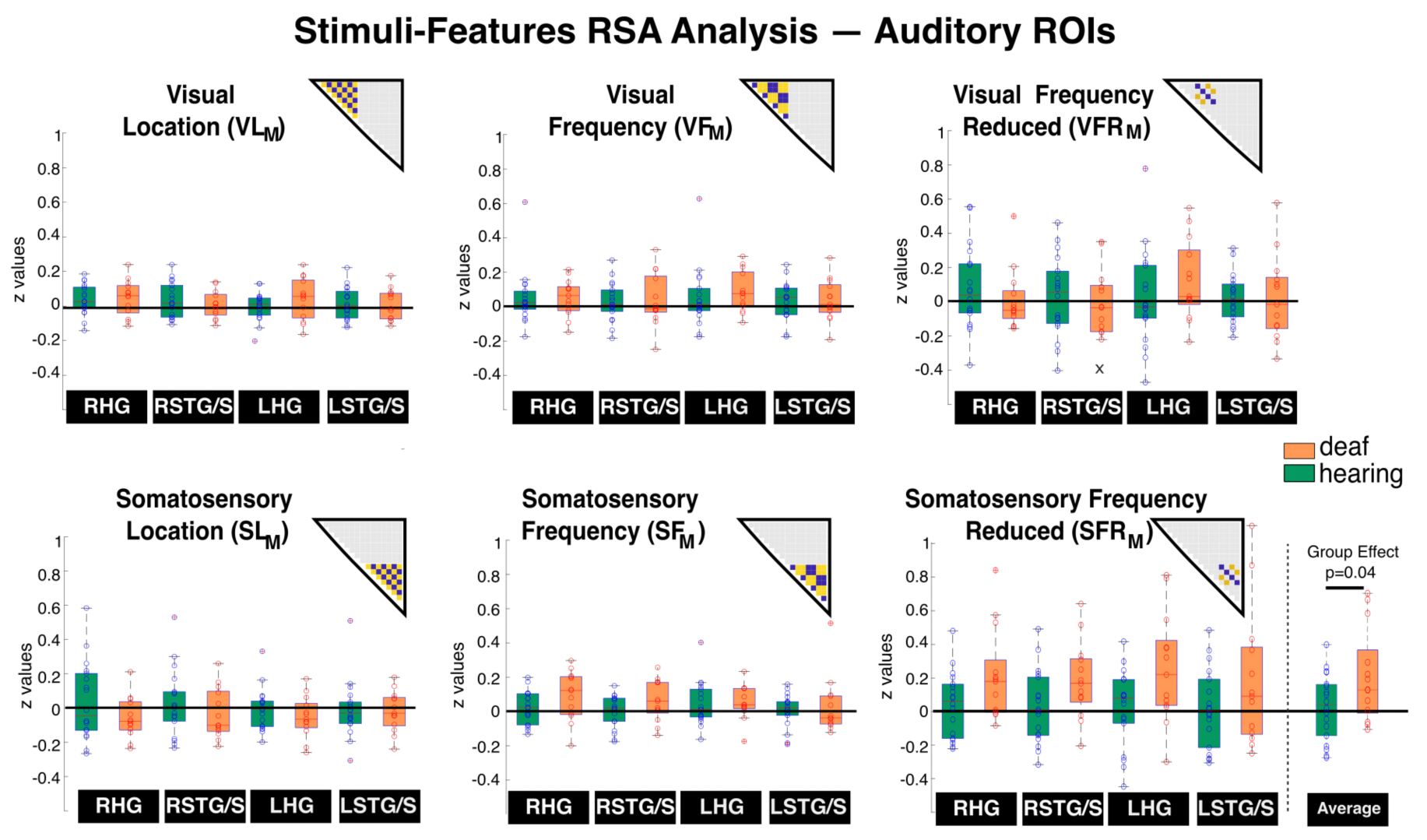
Representation of stimuli-features in auditory ROIs. The figure shows the Fisher-transformed z values for the Pearson correlation coefficients between the theoretical models for stimuli-features and each participant’s RDM. On each box, the edges indicate the 25^th^ and 75^th^ percentiles, and the black horizontal line inside the box indicates the median. RHG: right Heschl’s gyrus; LHG: left Heschl’s gyrus; RSTG/S: right superior temporal gyrus and sulcus; LSTG/S: left superior temporal gyrus and sulcus. VL**_M_**: Visual Location Model. VF**_M_**: Visual Frequency Model. VFR**_M_**: Visual Frequency Reduced Model. SL**_M_**: Somatosensory Location Model. SF**_M_**: Somatosensory Frequency Model. SFR**_M_**: Somatosensory Frequency Reduced Model. See also [VL_M_ ANOVA](https://osf.io/et2un/files/qpbhn), [VF_M_ ANOVA](https://osf.io/et2un/files/jya7v), [VFR_M_ ANOVA](https://osf.io/et2un/files/gytuk), [SL_M_ ANOVA](https://osf.io/et2un/files/pqc3z), [SF_M_ ANOVA](https://osf.io/et2un/files/9nxzh) and [SFR_M_ ANOVA](https://osf.io/et2un/files/pk85y).

There was also a significant interaction between ROI * Group in the VL_M_ (F(1,29)=4.47, p=0.04, ηp^2^=.13)), but none of the post-hoc t-tests showed significant differences between groups (all p >.1; see [VL_M_ ANOVA](https://osf.io/et2un/files/qpbhn); see t-tests in [Supp Mat 3 VL_M_](https://osf.io/et2un/files/usya5)).

In summary, these results show that the SFR_M_ z-scores were significantly higher than zero in the deaf group, and significantly different from those of the hearing group. The SFR_M_ tests if there is any frequency representation while the spatial location of the stimuli is kept constant. These results show that there is information about low-level stimuli features in the auditory cortex of deaf individuals. This was only the case for somatosensory frequency while spatial stimulation was kept constant.

##### Representations of task information (Fig 8)

We tested two models of task representations: the Task Modality-Dependent model (TMD_M_) and the Task Modality-Independent model (TMI_M_). Only in the TMD_M_ the ANOVA revealed group differences, with a significant ROI * Hemisphere * Group interaction (F(1,29)=5.34, p=.028, ηp^2^=.16) (see full ANOVA results [here](https://osf.io/et2un/files/pqx2s)). Post-hoc t-tests showed a significant difference between groups in RHG (t(29)=3.15, p=.004) (Fig. 8), reflecting a stronger fit in the deaf participants in this ROI. Marginal means for the z-scores of the TMD_M_ were significantly higher than zero in both groups (Deaf: mean=0.30, 95% CI=[0.17, 0.43], t(29)=5.6, p<.001; Hearing: mean=0.19, 95% CI=[0.06, 0.32], t(29)=3.5, p=.003; Fig 8)

**Figure 8.**
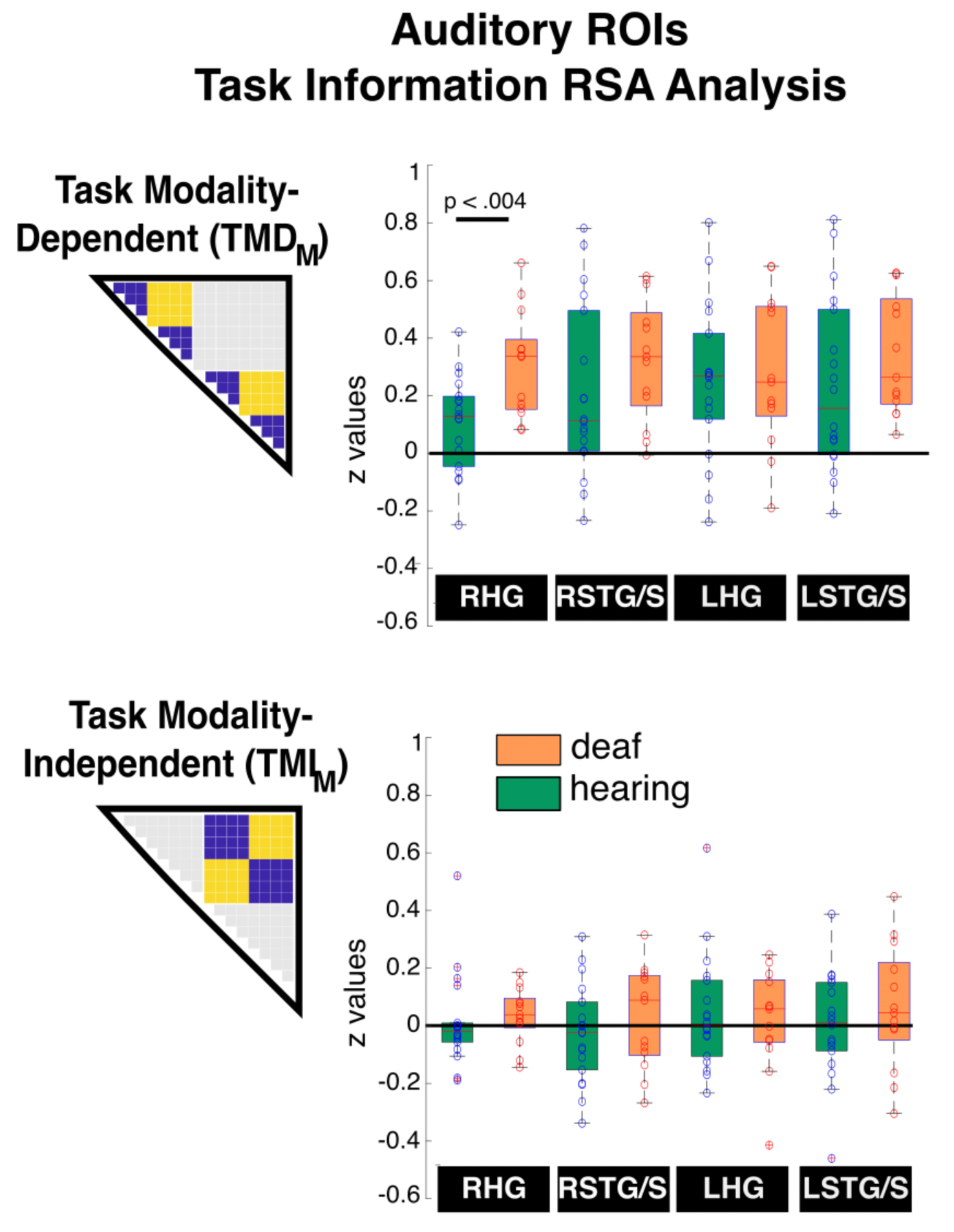
Representation of task information in auditory ROIs. The figure shows the Fisher-transformed z values for the Pearson correlation coefficients between the theoretical models and each participant’s RDM. On each box, the edges indicate the 25^th^ and 75^th^ percentiles, and the black horizontal line inside the box indicates the median. RHG: right Heschl’s gyrus; LHG: left Heschl’s gyrus; RSTG/S: right superior temporal gyrus and sulcus; LSTG/S: left superior temporal gyrus and sulcus. See also [TMD_M_ ANOVA](https://osf.io/et2un/files/pqx2s) and [TMI_M_ ANOVA](https://osf.io/et2un/files/kj7qb).

The z-scores of the TMI_M_ model were not significantly higher than zero in any of the groups (Deaf: mean=-0.035, 95% CI=[-0.11, 0.04], t(29)=-.96 p=0.34; Hearing: mean=0.017, 95% CI=[-0.06, 0.09], t(29)=0.47, p=.64), see full [TMI_M_ ANOVA results here](https://osf.io/et2un/files/kj7qb); Fig 8).

These results show that auditory areas code information about tasks in other sensory modalities, both in deaf and in hearing individuals. Our findings show that this representation of task information is enhanced in RHG as a consequence of deafness.

##### Representations of sensory modality (Fig 9)

In auditory ROIs, the ANOVA for the MOD_M_ showed a significant main effect of Group (F(1,29)=4.63, p=0.04, ηp^2^=.14), with overall higher correlations coefficients in the group of deaf individuals (Fig 9; [MOD_M_ ANOVA](https://osf.io/et2un/files/an7ps)). There was also a significant interaction Group * ROI (F(1,29)=4.62, p=.04, ηp^2^=.14). Independent samples t-tests showed a significant difference between groups in STG/S (t(29)=3.23, p=.003;), but not in HG (t(29)=1.18, p=.25; [Supp Mat 3 MOD_M_](https://osf.io/et2un/files/f2vxr)). Marginal means were significantly higher than zero in both groups (Deaf: mean=1.02, 95% CI=[0.82, 1.22], t(29)=11.8, p<.001; Hearing: mean=0.75, 95% CI=[0.55, 0.96], t(29)=8.72, p<.001).

**Figure 9.**
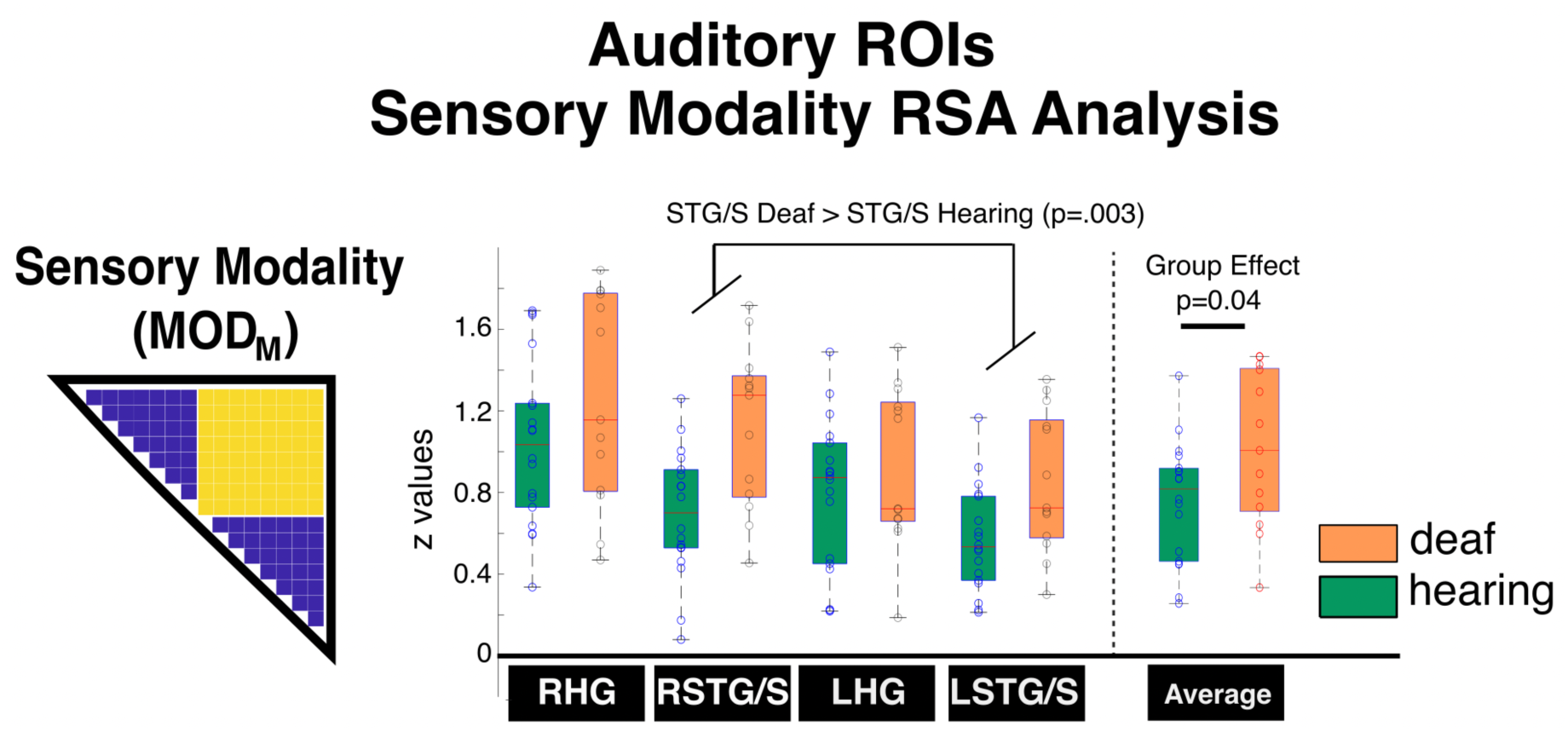
Representation of sensory modality information. The figure shows the Fisher-transformed z values for the Pearson correlation coefficients between the theoretical models and each participant’s RDM. On each box, the edges indicate the 25^th^ and 75^th^ percentiles, and the black horizontal line inside the box indicates the median. RHG: right Heschl’s gyrus; LHG: left Heschl’s gyrus; RSTG/S: right superior temporal gyrus and sulcus; LSTG/S: left superior temporal gyrus and sulcus. See also [MOD_M_ ANOVA](https://osf.io/et2un/files/an7ps).

These findings show that there is representation of modality information in auditory areas in both deaf and hearing individuals. A significant main effect of Group suggests that this representation is enhanced by sensory experience, and the significant interaction suggests that this enhancement is stronger in STG/S.

## DISCUSSION

Here we studied how sensory experience shapes the function of cortical sensory regions. Specifically, we investigated the role of the auditory cortex in implementing representations of sensory and task information across temporal and spatial dimensions. Our experiment revealed visual and somatosensory crossmodal plasticity in the auditory cortex of deaf individuals. Only deaf individuals showed somatosensory responses in HG, and visual and somatosensory responses in STG/S. The RSA analysis showed that the auditory cortex in deaf individuals represents information about task, sensory modality, and somatosensory frequency. Critically, task and modality representations were also found in the auditory cortex of hearing individuals, with significant differences between groups implying that these representations are enhanced as a consequence of deafness.

Our hypotheses were that: a) a representation of visual and somatosensory stimuli features in the auditory cortex of deaf individuals will support a functional preservation account; and b) representations of task and sensory modality in deaf individuals will support a functional change account. Instead, what we found was a more complex pattern that does not conform to this strict dichotomy. For task and sensory modality, we saw that both groups had significant representations of these dimensions. However, the functional relevance might be different, given that we did not observe a significant increase in the univariate response in hearing individuals. In contrast, for sensory features, the representation was different between groups. Yet, the underlying function appears analogous: somatosensory frequency processing in the auditory cortex of deaf individuals versus auditory frequency processing in hearing individuals. Overall, these findings suggest that crossmodal plasticity relies on representational and functional configurations that are present across individuals and modulated by sensory experience.

### Crossmodal Differences Between Groups in Auditory Regions

The univariate fMRI analysis showed the following crossmodal plasticity effects:

i. in HG, significant responses only to somatosensory stimulation in deaf individuals;
ii. in STG/S, significant differences between groups in both sensory modalities, with stronger responses for temporal tasks.

These findings suggest different profiles of crossmodal plasticity in primary (HG) and secondary auditory areas (STG/S).

In addition, the RSA analysis showed significant group differences for the Modality (MOD_M_) and Somatosensory Frequency Reduced (SFR_M_) models across all auditory ROIs. There was also a stronger representation of task information (Task Modality-Dependent Model, TMD_M_) for deaf individuals in RHG. Despite these group differences, it is striking that representations of task and sensory modality are significantly greater than zero in all auditory areas in both groups, suggesting that sensory cortices process this information in all individuals.

Overall, our findings show different univariate responses despite similar representations for task and modality in hearing and deaf individuals. Our results suggest that crossmodal plasticity impacts different levels of representation in the auditory cortex, including features, task and sensory modality. They also show that information about task and sensory modality represented in auditory regions is preserved across individuals, but modulated by sensory experience. To the best of our knowledge, this is the first study to address the representation of these multiple dimensions (stimuli features, task and sensory modality). Only a small number of studies in deaf individuals have used multivariate methods or model-based analyses of neural dynamics (Almeida et al. 2015; Kumar et al. 2024; Setti et al. 2023), and none has tested whether dimensions beyond sensory processing are represented in auditory cortex. Thus, our findings are fundamental for our understanding of crossmodal plasticity mechanisms and how sensory experience shapes the functional specialisation of brain regions, showing that auditory areas can represent different dimensions of information that go beyond sensory information.

### Similar Representation of Task and Modality Information Across Groups

Our study was designed to elucidate whether crossmodal plasticity effects in deaf individuals are due to the representation of information about sensory features, task and/or modality. Initially, we hypothesised that auditory cortex representations of task and modality information would suggest a functional change, expecting to observe these representations in deaf individuals, but not in hearing controls. What we found instead is that task and modality representations are found in both groups of deaf and hearing individuals. These findings suggest that crossmodal plasticity effects arise from representations that are present independently of sensory experience, rather than arising from the implementation of distinctly new functions in deaf individuals.

These findings agree with previous research showing that it is possible to decode different modalities of sensory stimulation from fMRI activity of an unstimulated primary sensory cortex. For instance, it is possible to decode whether a participant is receiving auditory or somatosensory stimulation from activity in unstimulated V1 (Liang et al., 2013). Critically, our results go beyond by showing modality responses in unstimulated cortices, and showing that sensory regions have distinct patterns of response for the task at hand, even when tasks are performed on sensory information other than their main sensory input.

While representations of task and modality were present in both groups in auditory areas, significant group differences show that these representations are enhanced as a consequence of deafness. The reduced auditory inputs experienced by deaf individuals may allow modality and task representations to become more functionally relevant. Previous findings have indeed suggested a functional relevance for the crossmodal plasticity effects found in univariate fMRI analysis (Manini et al. 2022), given the significant correlation between behavioural responses and the activation of the auditory cortex of deaf individuals during a visual switching task.

### Implications of Task and Modality Representations in Deaf and Hearing Individuals

What underlies the task and modality effects observed in deaf and hearing individuals? Research across several species has shown that putatively unimodal sensory cortices receive input from other sensory modalities (Bizley and King 2012; Meredith and Lomber, 2017; Kayser et al. 2010; Egea-Weiss et al. 2025; Wang et al. 2008). For example, the auditory cortex can integrate information about visual temporal dynamics, enhancing auditory object features that align with the temporal coherence of the visual stimuli (Bizley and King, 2012; Kayser et al. 2010). It is possible that these signals reach the auditory cortex also in deaf individuals, and that they are useful for cognition and perception in the absence of auditory inputs. For instance, Petro et al. (2017) suggest that crossmodal sensory signals are not always used for active perception. They suggest that these signals could be feedback mechanisms used for imagery and mind wandering, and while they many not be relevant for immediate perception, they can contribute to shaping future behaviour and cognition.

It has also been suggested that crossmodal signals in unstimulated cortices could be markers of conscious perception. Using at threshold sensory stimulation, Sanchez et al. (2020) showed that crossmodal signals can be decoded in unstimulated cortices, but only when they were consciously perceived. These findings show that these crossmodal signals are not automatic and require conscious awareness, and may reflect neural correlates of conscious perception.

Another possibility is that task and modality representations reflect attentional processing, which has been suggested as one of the roles of the reorganised auditory cortex (Cardin et al., 2023; Manini et al., 2022; Seymour et al., 2017). Using optical imaging, (Seymour et al., 2017) showed crossmodal plasticity effects in the right posterior temporal cortex during a visual selective attention paradigm. They also showed that the level of activity in this area correlated with behavioural performance in an independent off-line attention task across both groups of deaf and hearing individuals, suggesting a continuum in the response profile between deaf and hearing individuals. This posterior part of the auditory cortex is anatomically adjacent to the temporo-parietal junction (TPJ), an area that has a role in re-orientation of attention and in representing task-relevant information (Corbetta et al., 2008; Geng & Mangun, 2011). We have previously suggested that the function of the neighbouring TPJ could influence crossmodal plasticity effects (Cardin et al., 2023; Manini et al., 2022). The patterns of representations across tasks and modalities found here are consistent with this role.

In future studies, it would be important to determine if information beyond sensory processing is also represented for other cognitive dimensions in early sensory cortices. For example, in early blind individuals, areas of the visual cortex respond to spoken language stimulation (Bedny et al., 2011; Burton et al., 2002; Röder et al., 2002). Seydell-Greenwald et al. (2023) also showed univariate fMRI responses to spoken language in the visual cortices of sighted participants, which suggests a common function. Future research should investigate whether the representation of spoken language information is the same between sighted and blind individuals, and whether these representations are enhanced as a consequence of blindness. Our findings here suggest that such a shared mechanism may indeed exist.

### Plasticity Implications of Stimuli Features Representations

We found evidence of representation of somatosensory stimuli features in the auditory cortex of deaf individuals, but no evidence of representation of visual features.

There is a close interaction between auditory and somatosensory frequency processing in the brain. This is demonstrated by evidence of somatosensory responses in the auditory cortex of humans and animal models with full hearing (Caetano & Jousmäki, 2006; Foxe et al., 2002; Kayser et al., 2005; Nordmark et al., 2012), as well as by the representation of auditory frequency information in the somatosensory cortex (Pérez-Bellido et al., 2018).

This interaction provides a useful framework for interpreting our current results. In deaf individuals, our RSA findings suggest that the representation of somatosensory frequency occurs in a spatially-dependent way, rather than in an abstract frequency representation. We found no evidence of somatosensory frequency representation in auditory areas of hearing individuals. However, it is likely that the representation of somatosensory frequency in deaf individuals relies on the same mechanisms used to represent auditory temporal frequency in hearing individuals. In other words, the underlying computations might be the same (frequency processing), but the represented sensory information is different.

This processing of different sensory information, potentially relying on the same underlying functional mechanisms, has been previously been suggested for other stimuli and populations in crossmodal plasticity studies, including:

1. the visual cortices of blind individuals for the representation of categories, shapes, letters, objects and space conveyed by sound or touch (Amedi et al., 2007; Collignon et al., 2011; Mattioni et al., 2020; Pietrini et al., 2004; Striem-Amit et al., 2012; Striem-Amit & Amedi, 2014; van den Hurk et al., 2017; Xu et al., 2023).
2. the auditory cortex of deaf cats for the processing of visual location and motion (Lomber et al., 2010)
3. the auditory cortex of deaf individuals for sign language processing (Cardin et al., 2013, 2016; MacSweeney et al., 2002), faces (Benetti et al., 2017), and rhythmic sequences (Bola et al., 2017).

Contrary to the somatosensory findings, we found no evidence of a representation of visual features in auditory areas in deaf individuals. This suggests that, in the visual modality, the underlying representations resulting in stronger univariate responses in auditory areas in deaf individuals are not a result of feature processing. In other words, changes in the univariate responses observed in the visual condition in auditory regions of deaf individuals cannot be attributed to the representation of the physical properties of the stimuli. Instead, univariate changes are likely indicative of functions such as the specific set of cognitive computations that allow the successful completion of a task, or crossmodal attention effects.

### Differences in Effects Elicited by Visual and Somatosensory Stimulation

We found clear differences in the responses elicited by stimulation in the somatosensory and visual modalities. In HG we found strong univariate responses to the somatosensory tasks, but not the visual ones; while STG/S regions were responsive to both sensory modalities, with a stronger response for temporal tasks.

These findings confirm previous results from Karns et al., (2012), who showed stronger somatosensory, rather than visual, responses in HG. The recruitment of primary auditory cortex for somatosensation has also been shown in animal studies of early deafness (Hunt et al., 2006; Meredith & Allman, 2012; Meredith & Lomber, 2011). This difference in visual and somatosensory responses could represent potential different routes for crossmodal plasticity. While fMRI measurements do not allow us to determine the inputs that give rise to the observed responses, it is worth speculating whether visual and somatosensory stimuli have different pathways to the auditory cortex of deaf individuals. Evidence from animal models shows that deafness increases somatosensory responses and somatosensory projections to the cochlear nucleus (Shore et al., 2008; Zeng et al., 2012), which is the first auditory relay in the central nervous system. This increased somatosensory information could follow the auditory pathways and reach the primary auditory cortex, potentially resulting in the somatosensory responses that we see in HG in deaf individuals.

The pattern of responses for visual stimulation suggest a different explanation. Univariate visual responses in our experiment were only found in STG/S, and not in HG. This could be a result of visual signals arriving from cortico-cortical connections directly into the secondary auditory areas. Magnetoencephalography (MEG) studies of sign language and face processing support this notion. Leonard and colleagues (2012) investigated sign language processing in the left superior temporal cortex of deaf individuals. They examined whether activations occurred during an early sensory processing window (80-120 ms), or a later window (200-400 ms) associated with lexico-semantic processing. They found activity in the auditory cortex of deaf individuals peaking around 300-350ms, supporting the latter. In addition, Benetti et al. (2017) found face-specific responses in the right superior temporal cortex of deaf individuals, but not in hearing individuals. The peak for this face selectivity in the temporal cortex (‘deaf temporal face area’) is 192ms, 16ms after the peak in the Fusiform Face Area (FFA) at 176ms. None of these MEG studies found early responses coherent with early sensory processing in the auditory cortex of deaf individuals.

Overall, our findings showing modality-specific effects could reflect different pathways for inputs to the reorganised auditory cortex, but this will need to be addressed in further studies. While we found somatosensory frequency features represented in a spatially dependent manner in the auditory cortex of deaf individuals, there was no evidence of representations of visual sensory features, suggesting different modality constraints.

## Conclusion

Our study provides evidence of crossmodal plasticity in the auditory cortex of deaf individuals, highlighting its dual role in cognitive processing and sensory-specific representations. Our findings show that crossmodal task and modality representations are found in the auditory cortex of deaf and hearing individuals, and are enhanced in deaf individuals. This indicates that crossmodal plasticity relies on shared representations that are present in both deaf and hearing individuals. These results suggest that sensory areas can adapt their functional profile in response to sensory experience and enhance their processing of information from other modalities. They also suggest active roles for sensory areas in cognitive processes, beyond the representation of sensory features, which can serve as the basis for different functional destinies depending on the sensory experience of the individual.

## Acknowledgements

The authors would like to thank all the deaf and hearing participants who took part in the study, and Dr Elisa Infanti and Dr Davide Bono for their assistance with MRI data acquisition.

---

**Supplementary Materials 1.**
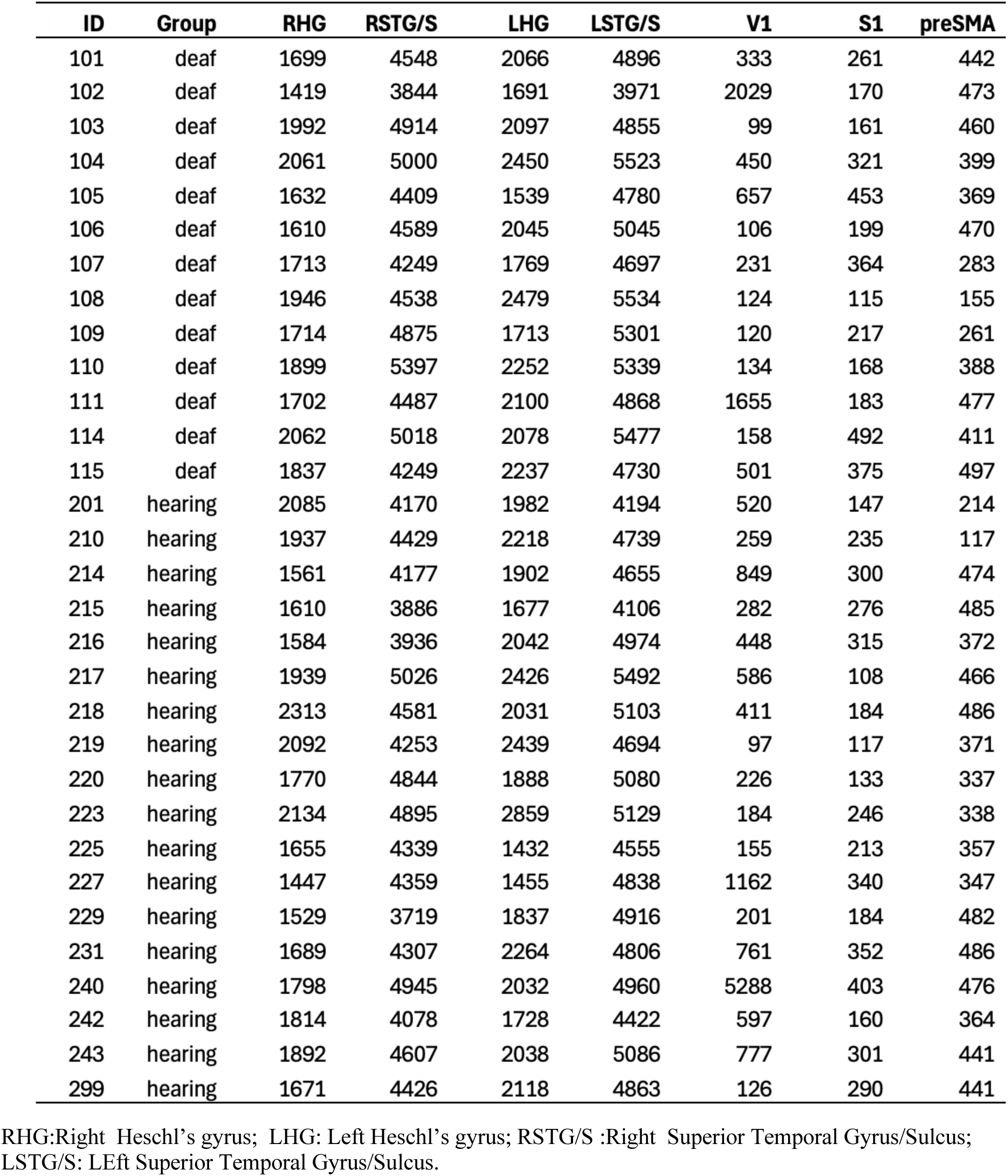
Number of voxels per ROI.

**Supplementary Materials 2.** Univariate Analysis Independent Samples T-tests: Auditory ROIs.

Group Differences in Right Heschl's Gyrus (Independent Samples T-Tests)
|  | t | df | p | Cohen's d | SE Cohen's d | 95% CI for Cohen's d |  |
| --- | --- | --- | --- | --- | --- | --- | --- |
|  |  |  |  |  |  | Lower | Upper |
| All | 3.455 | 29 | 0.002 | 1.258 | 0.440 | 0.466 | 2.031 |
| Somatosensory | 4.177 | 29 | < .001 | 1.520 | 0.471 | 0.697 | 2.323 |
| Visual | 0.758 | 29 | 0.455 | 0.276 | 0.368 | -0.443 | 0.990 |

*Note.* Student's t-test. All: average across all trials. Somatosensory: Average across somatosensory trials. Visual: Average across visual trials.

Group Differences in Left Heschl's Gyrus Independent Samples T-Test
|  | t | df | p | Cohen's d | SE Cohen's d | 95% CI for Cohen's d |  |
| --- | --- | --- | --- | --- | --- | --- | --- |
|  |  |  |  |  |  | Lower | Upper |
| All | 2.517 | 29 | 0.018 | 0.916 | 0.406 | 0.158 | 1.660 |
| Visual | -0.117 | 29 | 0.908 | -0.043 | 0.364 | -0.756 | 0.671 |
| Somatosensory | 3.519 | 29 | 0.001 | 1.281 | 0.442 | 0.486 | 2.057 |
| Temporal | 3.034 | 29 | 0.005 | 1.104 | 0.424 | 0.329 | 1.863 |
| Spatial | 1.673 | 29 | 0.105 | 0.609 | 0.383 | -0.126 | 1.334 |

*Note.* Student's t-test. All: average across all trials. Somatosensory: Average across somatosensory trials. Visual: Average across visual trials. Temporal: Average across temporal trials. Spatial: Average across spatial trials

Group Differences in Right Superior Temporal Gyrus/Sulcus (Independent Samples T-Test)
|  | t | df | p | Cohen's d | SE Cohen's d | 95% CI for Cohen's d |  |
| --- | --- | --- | --- | --- | --- | --- | --- |
|  |  |  |  |  |  | Lower | Upper |
| All | 4.372 | 29 | < .001 | 1.591 | 0.479 | 0.759 | 2.403 |
| Temporal | 5.127 | 29 | < .001 | 1.866 | 0.516 | 0.996 | 2.714 |
| Spatial | 3.023 | 29 | 0.005 | 1.100 | 0.423 | 0.325 | 1.859 |

*Note.* Student's t-test. All: average across all trials. Temporal: Average across temporal trials. Spatial: Average across spatial trials

Group Differences in Left Superior Temporal Gyrus/Sulcus (Independent Samples T-Test)
|  | t | df | p | Cohen's d | SE Cohen's d | 95% CI for Cohen's d |  |
| --- | --- | --- | --- | --- | --- | --- | --- |
|  |  |  |  |  |  | Lower | Upper |
| Temporal | 2.622 | 29 | 0.014 | 0.954 | 0.409 | 0.193 | 1.701 |
| Spatial | 0.874 | 29 | 0.389 | 0.318 | 0.369 | -0.403 | 1.034 |

*Note.* Student's t-test. Temporal: Average across temporal trials. Spatial: Average across spatial trials

**Supplementary Materials 3.** RSA Models Independent Samples T-tests: Auditory ROIs.

Independent Samples T-Tests Visual Location Model (VL<sub>M</sub>)
|  | t | df | p |
| --- | --- | --- | --- |
| Right HG | 0.257 | 29 | 0.799 |
| Right STG/S | -0.816 | 29 | 0.421 |
| Left HG | 1.176 | 29 | 0.249 |
| Left STG/S | -0.235 | 29 | 0.816 |

*Note.* HG: Heschl's gyrus; STG/S: Superior Temporal Gyrus/Sulcus

Independent Samples T-Tests Task Modality-Dependent Model (TMD<sub>M</sub>)
|  | t | df | p |
| --- | --- | --- | --- |
| Right HG | 3.147 | 29 | 0.004 |
| Right STG/S | 1.148 | 29 | 0.260 |
| Left HG | 0.328 | 29 | 0.746 |
| Left STG/S | 1.085 | 29 | 0.287 |

*Note.* HG: Heschl's gyrus; STG/S: superior temporal gyrus/sulcus

Independent Samples T-Tests Sensory Modality Model (MOD<sub>M</sub>)
|  | t | df | p |
| --- | --- | --- | --- |
| AvHG | 1.182 | 29 | 0.247 |
| AvSTG/S | 3.232 | 29 | 0.003 |

*Note.* AvHG: Average of Heschl's gyrus ROIs; AvSTG/S: Average of Superior Temporal Gyrus/Sulcus ROIs

**Supplementary Figure 1.**
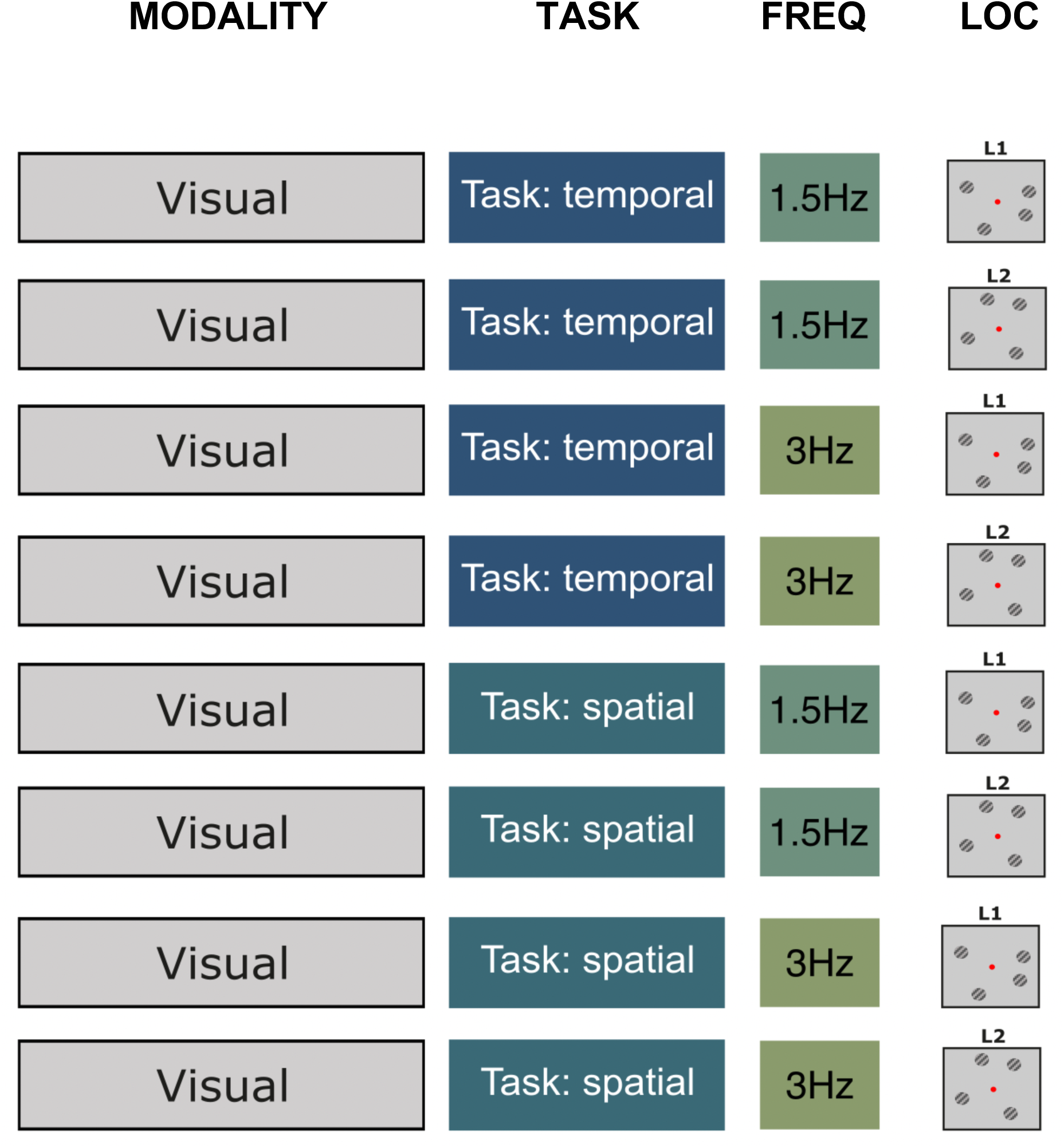
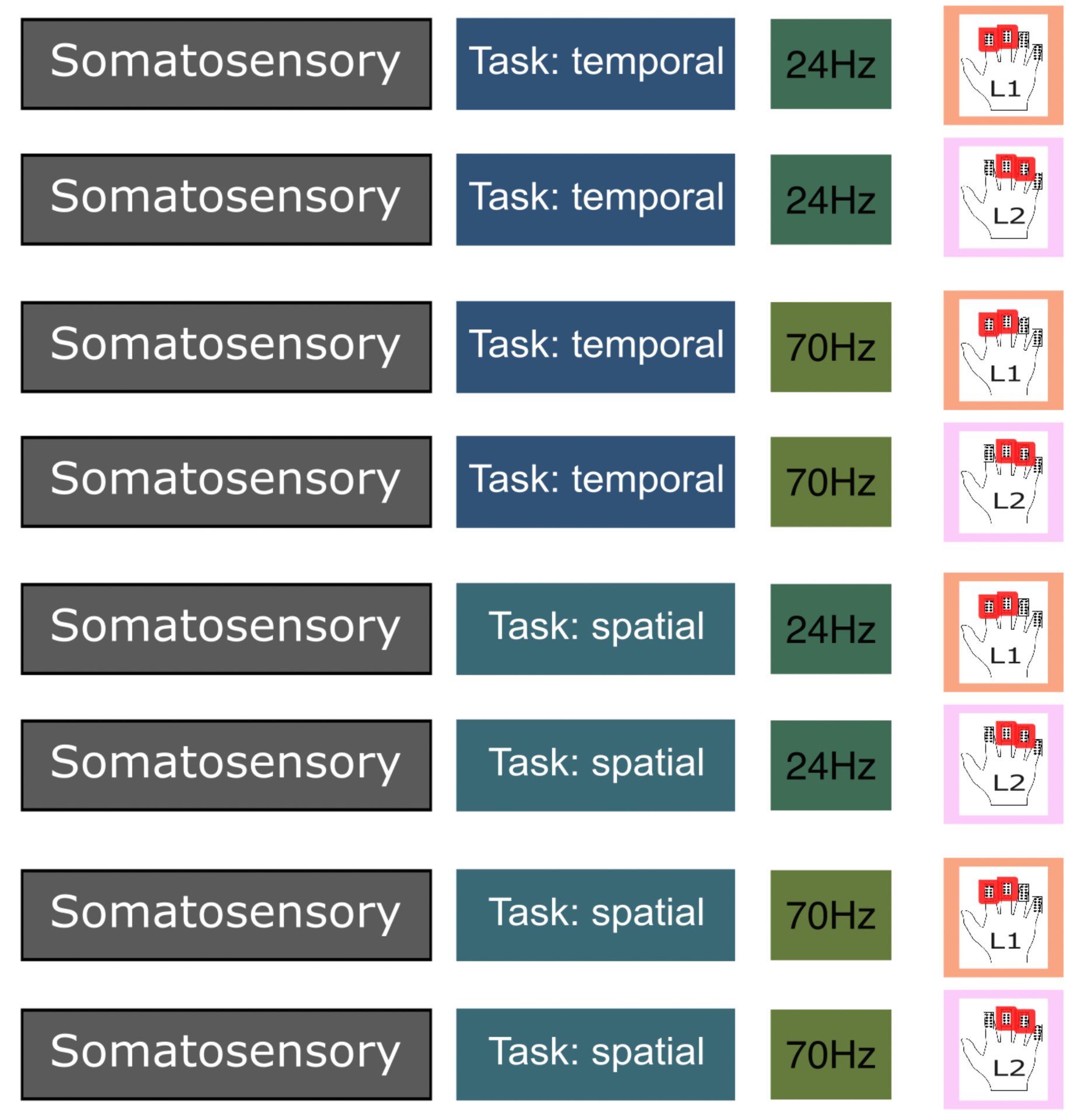
Examples of all visual and somatosensory samples. The order in the figure follows the order in the RSA matrices.

**Supplementary Figure 2.**
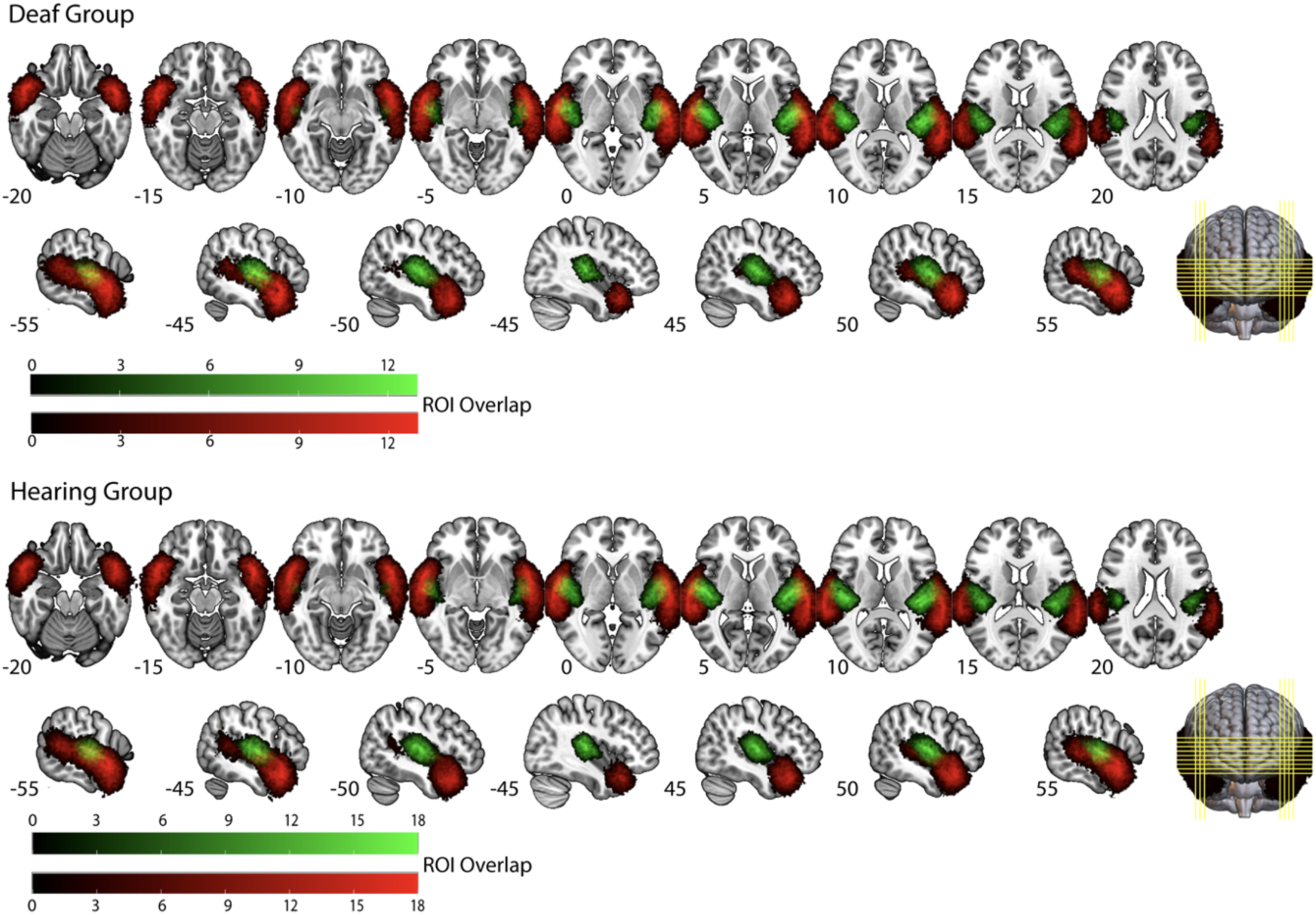
Auditory ROIs (group overlap). The figure shows the spatial overlap of auditory regions of interest (ROIs): Heschl’s Gyrus (HG, green) and Superior Temporal Gyrus/Sulcus (STG/S, red) for the deaf group (top panel) and the hearing group (bottom panel). Color intensity indicates the number of participants for whom a given voxel was included in the respective ROI (HG in green, STG/S in red).

**Supplementary Figure 3.**
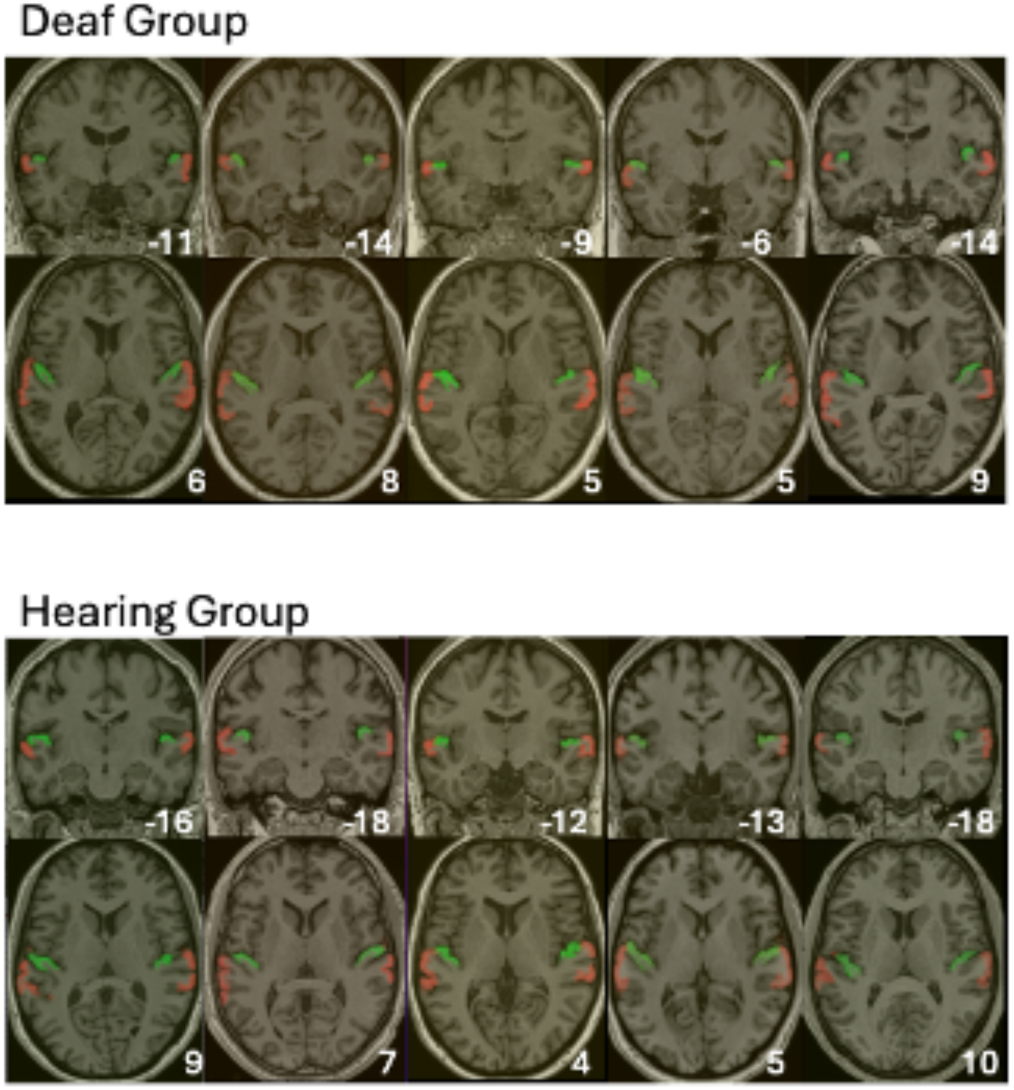
Auditory ROIs (individual examples). The figure shows examples of participant-specific auditory regions of interest (ROIs): Heschl’s Gyrus (HG, green) and Superior Temporal Gyrus/Sulcus (STG/S, red) from five deaf participants and five hearing participants. Numbers in white indicate the y (top rows) and z (bottom rows) coordinates.

